# Single cell analysis of blood mononuclear cells stimulated through CD3 and CD28 shows collateral activation of B and NK cells and demise of monocytes

**DOI:** 10.1101/2020.12.23.424147

**Authors:** Nathan Lawlor, Djamel Nehar-Belaid, Jessica D.S. Grassmann, Marlon Stoeckius, Peter Smibert, Michael L. Stitzel, Virginia Pascual, Jacques Banchereau, Adam Williams, Duygu Ucar

**Affiliations:** The Jackson Laboratory for Genomic Medicine, Farmington, CT, 06030, USA; 10X Genomics, Stockholm, Sweden; New York Genome Center, New York, NY 10013, USA; Institute of Systems Genomics, University of Connecticut, Farmington, CT 06032, USA; Department of Genetics and Genome Sciences, University of Connecticut, Farmington, CT 06032, USA; Ronay Menschel Professor of Pediatrics, Drukier Institute, Weill Cornell Medicine, New York, NY 10065, USA

**Keywords:** immune responses, single cell profiling, immune cell activation, LPS, antiCD3/CD28

## Abstract

Immune cell activation assays have been widely used for immune monitoring and for understanding disease mechanisms. However, these assays are typically limited in scope. A holistic study of circulating immune cell responses to different activators is lacking. Here we developed a cost-effective high-throughput multiplexed single-cell RNA-seq combined with epitope tagging (CITE-seq) to determine how classic activators of T cells (anti-CD3 coupled with anti-CD28) or monocytes (LPS) alter the cell composition and transcriptional profiles of peripheral blood mononuclear cells (PBMCs) from healthy human donors. Anti-CD3/CD28 treatment activated all classes of lymphocytes either directly (T cells) or indirectly (B and NK cells) but reduced monocyte numbers. Activated T and NK cells expressed senescence and effector molecules, whereas activated B cells transcriptionally resembled autoimmune disease- or age-associated B cells (e.g., CD11c, T-bet). In contrast, LPS specifically targeted monocytes and induced two main states: early activation characterized by the expression of chemoattractants and a later pro-inflammatory state characterized by expression of effector molecules. These data provide a foundation for future immune activation studies with single cell technologies (https://czi-pbmc-cite-seq.jax.org/).

**Graphical abstract:** 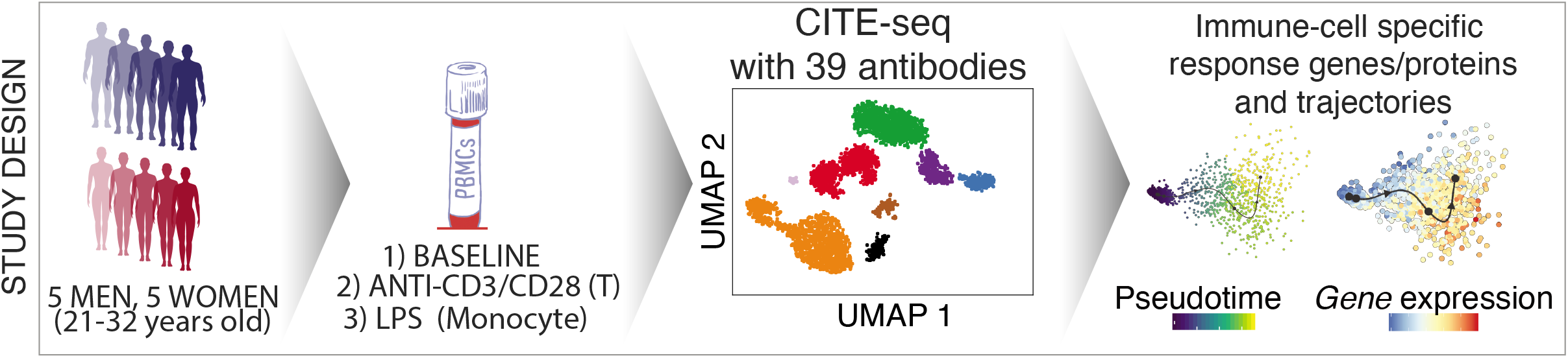

## Introduction

The immune system plays a central role not only in fighting infections, but also in the pathogenesis of many diseases. While access to disease-relevant tissues may be limited, blood profiling provides a minimally invasive method to assess immune function and health (1,2). Blood serves as a conduit for transport of immune cells throughout the body and circulating immune cells frequently display signatures of disease or immune dysfunction (i.e., immunosenescence) (3–5). Transcriptional profiling of blood provides an unbiased method for immune monitoring and has proved invaluable in uncovering novel disease mechanisms. *In vitro* activation of immune cells is an effective way to assess their functional capacity. For example, T cells can be activated with non-specific mitogens such as PHA (phytohemagglutinin) and ConA (Concanavalin A) (6), or more specifically through the T cell receptor and co-stimulatory receptors, either by crosslinking with antibodies against CD3 and CD28 (7), or with antigen-laden antigen presenting cells (8). Similarly, innate cells, including monocytes, can be activated through pattern recognition receptors (PRRs) with pathogen-associated molecular patterns (PAMPs) such as lipopolysaccharide (LPS) (9). Studying such cellular responses is an effective strategy to uncover functional defects and/or disease-associated phenotypes in human immune cells, and has been widely applied to autoimmune diseases (10–12), immunodeficiency, aging (13–16), and allergy (17).

Advances in single cell technologies can precisely uncover donor-, cell type-, and cell state-specific variation in immune cell functions from clinical samples containing mixed cell populations (18,19). Hence, these technologies are increasingly used to study *ex vivo* and *in vitro* cellular responses of diverse immune cell types (CD8^+^ T cells (20,21), dendritic cells (DCs) (22,23), and B cells (24)) as well as activated cells in immune diseases (25). However, single cell gene expression profiling of activated cells has unique data generation and analysis challenges including identifying and removing multiplet cells that increase in number as cellular interactions increase upon activation. Furthermore, avoiding batch effects is essential to distinguish activation-specific gene expression patterns from batch-induced changes. Another major limitation is the high cost of the methodology which prevents large scale studies. Recent advances can address some of these challenges. For example, multiplexing *via* hashtag oligonucleotides (26) and individuals’ genotypes (27) permits simultaneous sequencing of libraries from multiple individuals and conditions to reduce both the cost and technical batch effects. Finally, cellular indexing of transcriptomes and epitopes by sequencing (CITE-seq) technology (26) profiles epitopes (i.e., cell surface proteins) and gene expression levels from the same single cells and is particularly suitable for studying distinct immune cell types and states (e.g., activated and resting) and for filtering out multiplets among activated cells.

We combined cell hashing and genotype-based multiplexing to generate CITE-seq data for the deep analysis of circulating human immune cell responses to classic activators of adaptive (anti-CD3/CD28) and innate (LPS) immune cells. Responses to these activators have been studied over decades (6) *via* bulk profiling of sorted cells in a reductionist manner, which missed indirect effects of these activators on other cell types as well as the heterogeneity in responses at the single cell level. PBMCs from 10 healthy young donors were profiled at baseline and following activation *via* 1) anti-CD3 + anti-CD28 for selective T cell activation, or 2) LPS for monocyte activation using CITE-seq to quantify changes in both transcript and protein levels using 39 antibodies (e.g., CD3, CD4, CD14, CD56, CD69, CD25). Data from this study uncovered cell-specific responses to each condition at both the gene and protein level and uncovered direct and indirect effects of these activators on immune cell transcriptomes. This study will guide future single cell studies of activated immune cells in health and disease (**https://czi-pbmc-cite-seq.jax.org/**).

## Results

### Simultaneous protein and gene expression profiling of resting and activated PBMCs

PBMCs were isolated from the blood of 10 healthy young individuals (5 men, 5 women; 21-32 years old; Caucasian), and were cultured for 24 hours with LPS (10ng/ml), anti-CD3/CD28 (2μg/ml each) (Methods), or in medium alone (referred to as ‘Baseline’). Flow cytometry analysis confirmed that anti-CD3/CD28 treatment induced the expression of activation markers (CD25, CD69) on T cells (Fig. 1A, B), whereas LPS activated monocytes as shown by CD80 induction (Fig. 1C, D). To study the effects of these activation conditions at the single cell level, we pooled baseline and activated cells from the 10 individuals (30 samples in total) and used CITE-seq to simultaneously profile changes in levels of transcripts and 39 cell surface proteins (Table S1). To reduce experimental batch effects and costs, equal numbers of cells from all donors and conditions were pooled and sequenced together. Cells from different conditions were labeled with Cell Hashing Antibodies (BioLegend) (Fig. 1E), where three hashtag oligonucleotides (HTOs) were used to demultiplex data from each experimental condition (baseline, LPS, anti-CD3/CD28) using a Gaussian mixture model on HTO counts (Fig. S1A). Finally, donor genotypes were used to demultiplex data from each individual with *Demuxlet* (27) (Fig. 1F). Overall, 195,480 droplets were captured from all samples (Table S2).

**Fig. 1:**
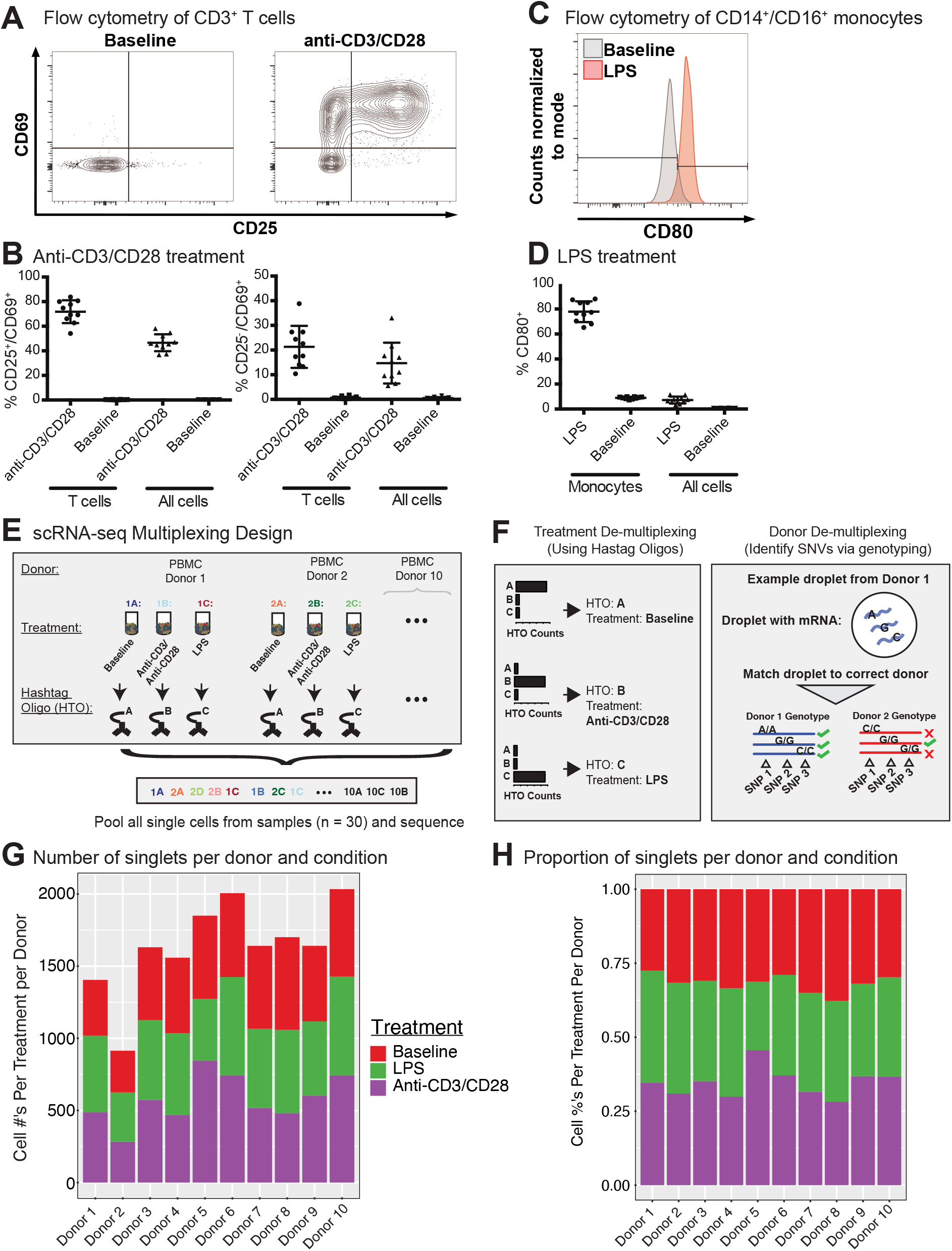
CITE-seq data before and after immune cell activation uncover distinct cell types and cell states. (A) Fluorescence-activated cell sorting (FACS) and enrichment of activated T cells at baseline and anti-CD3/CD28 conditions. T cells were gated as CD3^+^ T cells. (B) Flow cytometry data quantifying the proportion of activated cells (CD25^+^ CD69^+^ or CD25^-^ CD69^+^) after anti-CD3/CD28 stimulation. All cells = number of activated T cells as a percentage of all PBMCs; T cells = numbers of activated T cells as a percentage of all CD3^+^ T cells. (C) Distributions of CD80 expression in sorted monocytes and all PBMCs at baseline and LPS conditions. (D) Flow cytometry proportions of activated monocytes (CD80^+^) before and after LPS stimulation. All cells = number of activated monocytes as a percentage of all PBMCs; Monocytes = number of activated monocytes as a percentage of CD14^+^CD16^+^ monocytes. (E) Sample multiplexing was performed using cell hashtag oligonucleotide (HTO) antibodies, where a different HTO barcode was used for a specific treatment condition (e.g., HTO A for cells under baseline condition). (F) De-multiplexing of treatment information using HTOs (left), and donor of origin information using genetic variation across donors in the form of single nucleotide variations (SNVs)(right). (G) Numbers and (H) proportions of singlet cells retained per donor per condition after removing multiplet cells.

Cell multiplets from different sources were detected and eliminated with the help of multiplexing strategies (HTOs and genotypes) and the levels of cell-specific marker proteins (Fig. S1B, Methods). HTOs uncovered multiplets from different treatment conditions, which constituted ~13% (26,084/195,480) of total droplets in addition to 18,871(~9%) empty droplets (i.e., no HTO detected) (Fig. S1C). Using genotypes, we identified and filtered out multiplets from different individuals as well as cells that could not be assigned to an individual with high confidence (i.e., ambiguous) (Fig. S1D), resulting in 55,353 single cells with a mean of 1,932 unique molecular identifiers (UMIs) per cell. Empty droplets from cell hashing and ambiguous cells from *Demuxlet* overlapped extensively (83%, Fig. S1E). Finally, expression levels of cell type-specific proteins uncovered multiplets from different cell types (Methods). After multiplet removal and demultiplexing, an average of 1,638 (range 914-2034) cells per donor were detected (16,382 total cells) (Fig. 1G), with 855 genes detected per cell on average. Donors showed similar cell frequency distributions across conditions (Fig. 1H). The proportions of CD25^+^/CD69^+^ T cells from CITE-seq and flow cytometry data were highly correlated (Pearson’s R=0.96) (Fig. S1F) across individuals upon anti-CD3/CD28 stimulation, indicating that CITE-seq is an effective technology to capture and study immune cell activation.

### Anti-CD3/CD28 treatment activates all classes of lymphocytes

Baseline and anti-CD3/CD28*-*activated PBMCs were jointly analyzed using the top 500 most variably expressed genes and proteins (Fig. 2A; Methods). Cell clusters were annotated using the expression of marker proteins: CD19 and CD20 for B cells, CD16 and CD56 for NK cells, CD14 and CD11c for CD14^+^ monocytes, CD3 and CD4 for CD4^+^ T cells, CD3 and CD8a for CD8^+^ T cells, and CD45RA and CD45RO for naïve and memory T cells, respectively (Fig. 2B, S2A). Accordingly, baseline PBMCs from 10 individuals consisted of 60-82% T cells, 12-25% CD14^+^ monocytes, 4-15% NK cells, and 3-6% B cells (Fig. 2C), which was highly correlated with flow cytometry data from this cohort (Fig. S2B; Pearson’s R = 0.97). CD8^+^ memory T cells were the most variable cell type between individuals with a coefficient of variation (CV) of 0.65, followed by NK cells (CV=0.51).

**Fig. 2:**
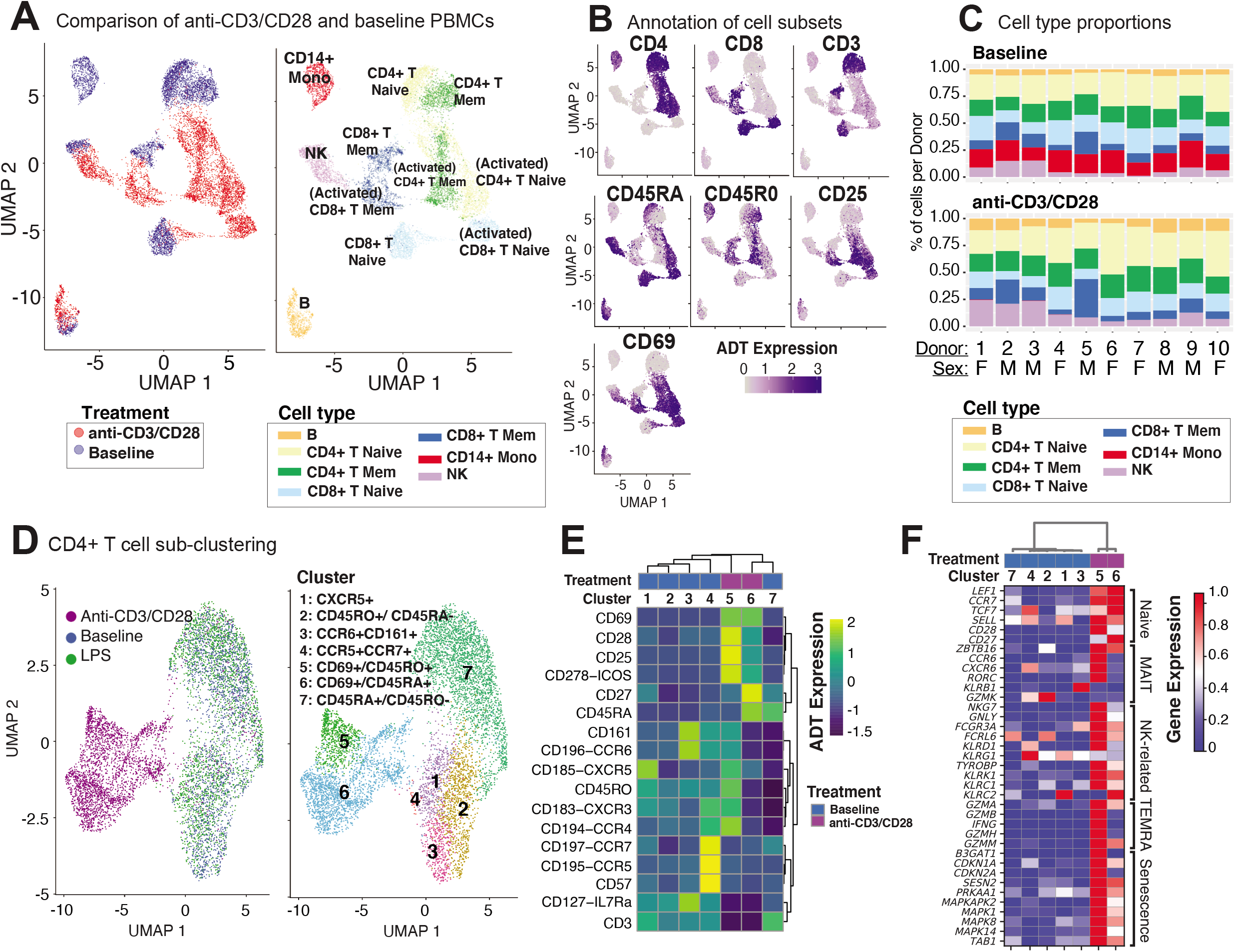
Anti-CD3/CD28 stimulation activates all classes of lymphocytes. (A) Clustering of PBMCs at baseline and anti-CD3/CD28 conditions separates lymphocytes (NK, T, and B cells) primarily by cell type and secondarily by condition. Cells are color-coded based on activation condition (left) and cell types inferred *via* CITE-seq antibodies (right). (B) Annotation of cells at baseline was performed using protein expression levels of CD3, CD4, CD8, CD45RA, CD45RO, and additional annotation of activated T cells was done using expression of CD25 and CD69. (C) Cell type proportions across 10 donors at baseline (top) and anti CD3/CD28 condition (bottom). Note the decline in CD14^+^ monocytes (red bars) upon anti-CD3/CD28 condition. (D) Unsupervised clustering of all CD4^+^ T cells separates anti-CD3/CD28-stimulated cells from those at baseline and LPS activation. Dimension reduction plot with CD4^+^ T cells colored by their resultant cell type identity. (E) Average scaled expression of protein markers used to identify and annotate CD4^+^ T cell subsets. (F) Average scaled expression of marker genes associated with naive, MAIT, NK, TEMRA, and senescence functions.

Anti-CD3/CD28 treatment activated all lymphocytes (Fig. 2A, left). PBMCs grouped primarily by cell type and secondarily by cell state (i.e., resting vs. activated) based on their annotations with cell surface markers (Fig. 2B, S2A). Labeled antibodies allowed the detection of naive (CD45RA^high^) and memory (CD45RO^high^) T cells. Activated T cells showed early (CD69) and late (CD25) activation markers and reduced CD3 expression (Fig. 2B). Indirect activation of NK and B cells was revealed by the induction of CD69 mRNA and protein (Fig. 2B). Notably, this treatment caused a significant decline in monocyte numbers and their percentages in PBMCs (Fig. 2C). This is likely due to the induction of apoptosis in monocytes *via* activated T cells (28)’(29) as apoptosis-associated genes were upregulated (e.g., *CTSL, TRAF1, TNF,* and *GZMB)* among T cell-monocyte multiplets detected in the anti-CD3/CD28 condition.

### Senescence and cytotoxicity molecules are upregulated in activated CD4^+^ T cells

Unsupervised clustering of CD4^+^ T cells using both gene and protein expression levels revealed 7 subgroups: 5 found at baseline and 2 additional ones following activation with anti-CD3/CD28 (Fig. 2D). CD4^+^ cells did not respond to LPS activation and clustered with baseline cells. At baseline, we detected CD45RA^+^CD45RO^-^ (cluster 7; naïve cells) and CD45RA^-^CD45RO^+^ (cluster 2; memory cells) subsets as well as less abundant CD45RO^+^CCR6^+^CD161^+^ (cluster 3; memory cells including Th17 cells), CD45RO^+^CXCR5^+^ (cluster 1; memory cells including Tfh cells), and CD45RO^+^CD57^+^CCR5^+^CCR7 cells ^+^ (cluster 4; other memory cells). CD45RO^+^CXCR5^+^ cells, which constituted ~8% of baseline CD4^+^ T cells (3-14% between donors) (Fig. S2C) were in an apparent resting state as they lacked ICOS (CD278) and CD69, in alignment with our previous observations (30). Cluster 3 cells (Fig. 2D, 2E) expressed chemokine receptor CCR6 (indicating tissue-specific homing properties) and C-type lectin receptor CD161 *(KLRB1)* and CD127 *(IL7R)* on their surface constituted ~9.9% of CD4^+^ T cells (Fig. S2C). Finally, we identified a smaller subset of memory cells that highly expressed CD57, CCR5, and CCR7 on their surface in addition to moderately expressing CXCR3 and CCR4 (Fig. 2E) and include Th1 (CCR5 and CXCR3) and Th2 (CCR3, CCR4) subsets (31). Activated CD4^+^ T cells clustered into two groups: cluster 5 (memory activated: CD69^+^ CD45RO^+^) and cluster 6 (naïve activated: CD69^+^ CD45RA^+^). Cells in both clusters expressed co-stimulatory proteins (most notably CD28, CD278) and activation marker proteins (CD69, CD25) (Fig. 2D). However, the expression of certain markers decreased upon activation (e.g., CD161, CCR6) limiting our ability to discriminate subsets among activated CD4^+^ T cells. T cell senescence genes (e.g., *SESN2, PRKAA1, MAPK1, TAB1)* were induced both in naive and memory CD4^+^ T cells upon activation (Fig. 2F). Moreover, activated memory cells (CD45RO^+^CD45RA^-^CD69^+^) highly expressed genes associated with terminally differentiated effector cells (T cells re-expressing CD45RA (TEMRA))(32), including cytotoxic molecules (e.g., GZMA, GZMB, IFNG, GZMH, GNLY) (Fig. 2F) (Table S3).

### Naive T cell responses to anti-CD3/CD28 stimulation are stronger than those of their memory counterparts

CD8^+^ T cells were composed of 6 distinct clusters: 4 found at baseline and 2 additional ones upon anti-CD3/CD28 activation. LPS did not activate CD8^+^ T cells as cells treated with LPS clustered together with baseline cells (Fig. 3A). Baseline clusters included: CD45RA^+^CD45RO^-^ (cluster 4; naïve), CD45RO^+^CD45RA^-^ (cluster 1; memory) cells, CD57^+^CXCR5^+^CD45RO^high^ cells (cluster 2; effector memory cells, ~8.6% of CD8+ T cells) and CD161^+^CCR6^+^ cells (cluster 5; MAIT, mucosal-associated invariant T cells, ~7.9% of CD8+ T cells) (Fig. 3B, Fig. S2C). Anti-CD3/CD28 activation elicited a strong response in CD8^+^ T cell subsets (Fig. 3A), resulting in activated memory cells (cluster 3; CD69^+^CD45RO^+^) and activated naïve cells (cluster 6; CD69^+^CD45RA^+^). Both activated clusters expressed high levels of activation proteins (CD25, ICOS, CD69) (Fig. 3B). When compared to activated memory cells, activated naïve cells expressed higher levels of the CD27 and CD28 proteins (Fig. 3B). Cluster 5 cells expressed marker genes of MAIT cells *(RORC, KLRB1,* and *CCR6* (33)) (Fig. 3C). Activated cells from clusters 3 and 6 expressed high levels of both cytotoxic (e.g., *GZMA, GZMK)* and senescence genes *(SESN2, CDKN1A, CDKN2A)* (Fig. 3C).

**Fig. 3:**
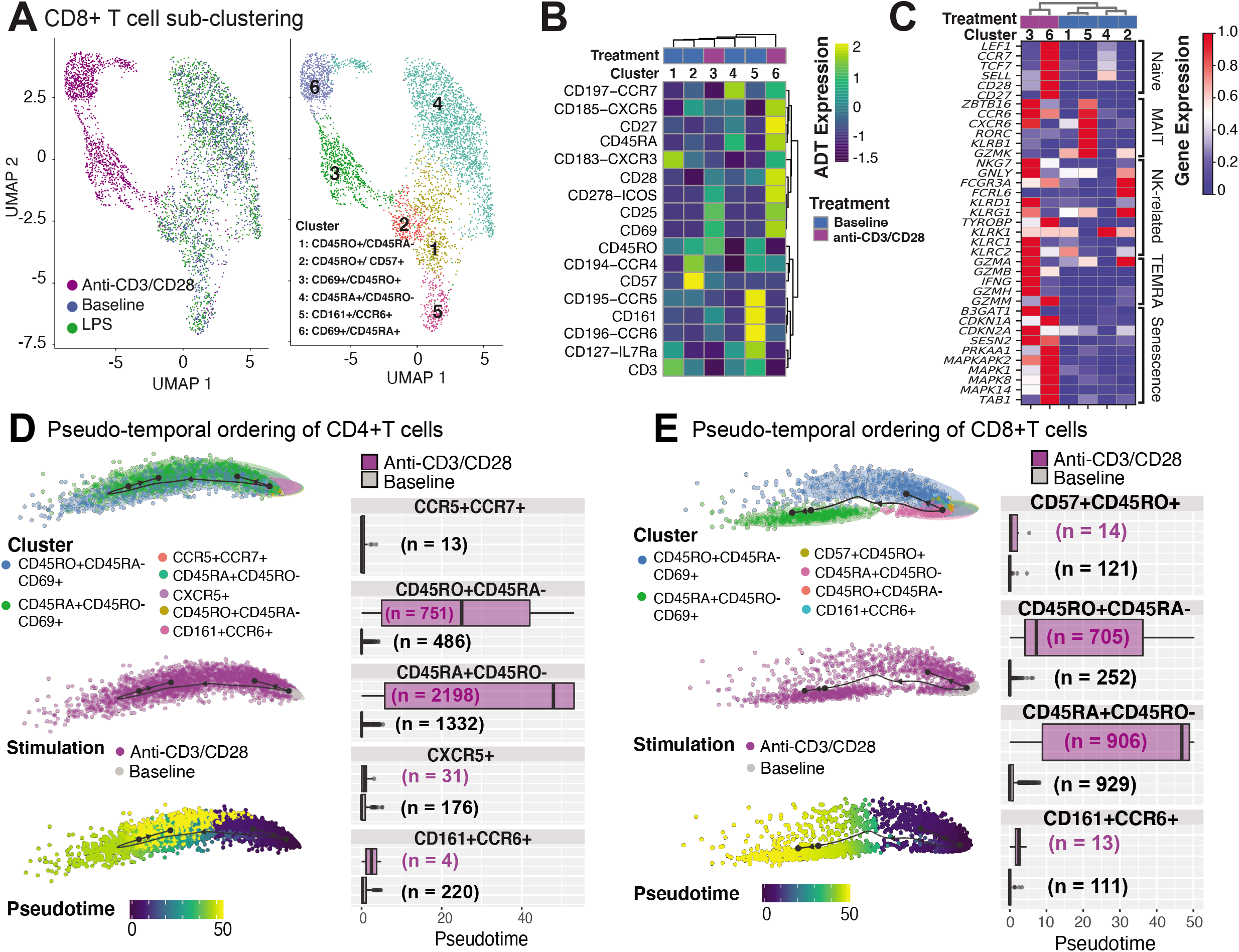
Annotation of CD8^+^T cell subsets and trajectory inference. (A) Unsupervised clustering of CD8^+^ T cells from all conditions reveals a separation of anti-CD3/CD28-stimulated cells from all others. Dimension reduction plot with CD8^+^ T cells colored by their resultant cell-type identity. (B) Average scaled expression of protein markers used to also identify CD8^+^ T cell subpopulations. (C) Average scaled expression of marker genes associated with naïve, MAIT, NK, TEMRA, and senescence functions. Pseudo-temporal ordering of (D) CD4^+^ and (E) CD8^+^ T cells at baseline and anti-CD3/CD28 conditions. Boxplots show the distributions of pseudotime states for each of the baseline/activated T cell subsets.

Differential expression analyses between activated and resting T cells revealed that 950 genes were upregulated in naïve and memory CD4^+^ and CD8^+^ T cells (Fig. S2D), including *IFNG,* Type I interferon-inducible genes *(ISG15, ISG20),* and interferon regulatory factors *(IRF1, IRF4)* (Table S4). Stimulation induced more genes in naïve (CD45RA^+^CD45RO^-^) T cells than memory (CD45RO^+^CD45RA^-^) T cells; 824 transcripts were induced specifically in naïve CD4^+^ and CD8^+^ T cells, while 494 were induced in CD8^+^ naive T cells (Fig. S2D). Naïve cell-specific response genes were enriched for KEGG pathways associated with cell cycle (e.g., *MCM2/3/5/6, CDKN2D, SMC1A, CHEK1),* carbon metabolism (e.g., *ESD, CS, SDHD, GPI),* and basal transcription factors (e.g., *ETF1, EIF4E, EIF4H, EIF4G1),* emphasizing the proliferative and expansive capacity of naive T cells upon activation (Table S4). Pseudo-temporal ordering of baseline and activated T cells using the Slingshot algorithm (34) revealed mostly linear trajectories for activation despite the existence of distinct memory (CD45RO^+^CD45RA^-^) subsets at baseline, thereby indicating that these different subsets respond similarly to anti-CD3/CD28 activation. Distinct subsets (e.g., CD161^+^CCR6^+^) were no longer detectable upon activation - potentially due to internalization of cell surface proteins - preventing further interrogation of their specific responses. Instead, our analyses revealed that, for both CD4^+^ T and CD8^+^ T compartments, activated naïve (CD45RA^+^CD45RO^-^CD69^+^) cells were ordered towards the end of the activation pseudotime trajectories, due to the greater transcriptomic changes in naive T cells upon anti-CD3/CD28 stimulation compared to memory cells (Figs. 3D,E). Taken together these data show that anti-CD3/CD28 activation has a significant effect on T cell transcriptomes, where naïve T cells go through more significant changes compared to their memory counterparts.

### Anti-CD3/CD28 activation of PBMCs result in indirect activation of NK cells

Anti-CD3/CD28 treatment induced CD69 protein expression on NK cells, and transcription of 74 genes (Table S5), 60 of which were shared with T cells, including cytotoxic molecules (e.g., *GZMB)* as well as interferon response genes (Fig. S2D, Table S4). Clustering analyses of NK and T cells, at baseline and after anti-CD3/CD28 activation, uncovered the transcriptional similarity between NK and CD8^+^ T cells (Fig. 4A). Interestingly, activated CD8^+^ T cells clustered distinctly from NK cells likely due to the upregulation of distinct molecules in T cells (Fig. S2D). To further characterize these cells, we derived MAIT, cytotoxicity, and senescence scores using relevant gene sets (Methods, Table S3). As expected, CD8^+^CD161^+^CCR6^+^ cells had the highest MAIT score. Activated NK cells (CD56^+^CD69^+^) showed the highest cytotoxicity score among all studied cell types, followed by activated CD8^+^ memory cells (CD8^+^CD45RO^+^CD69^+^) T cells (Fig. 4B). In alignment with this, upon activation, important cytotoxic molecules were up-regulated in NK cells, including perforin *(PRF1),* granulysin *(GNLY),* and granzymes *(GZMB)* (Table S5). In contrast, activated T cells had high senescence scores, particularly for activated naive CD8^+^ T cells (CD45RA^+^CD69^+^), which was in alignment with the increased expression of CDKN1A (encoding p21 protein) and CDKN2A (encoding p16 protein) along with other senescence-associated molecules (Fig. 3C). In contrast, baseline and activated NK cells displayed low senescence scores.

**Fig. 4:**
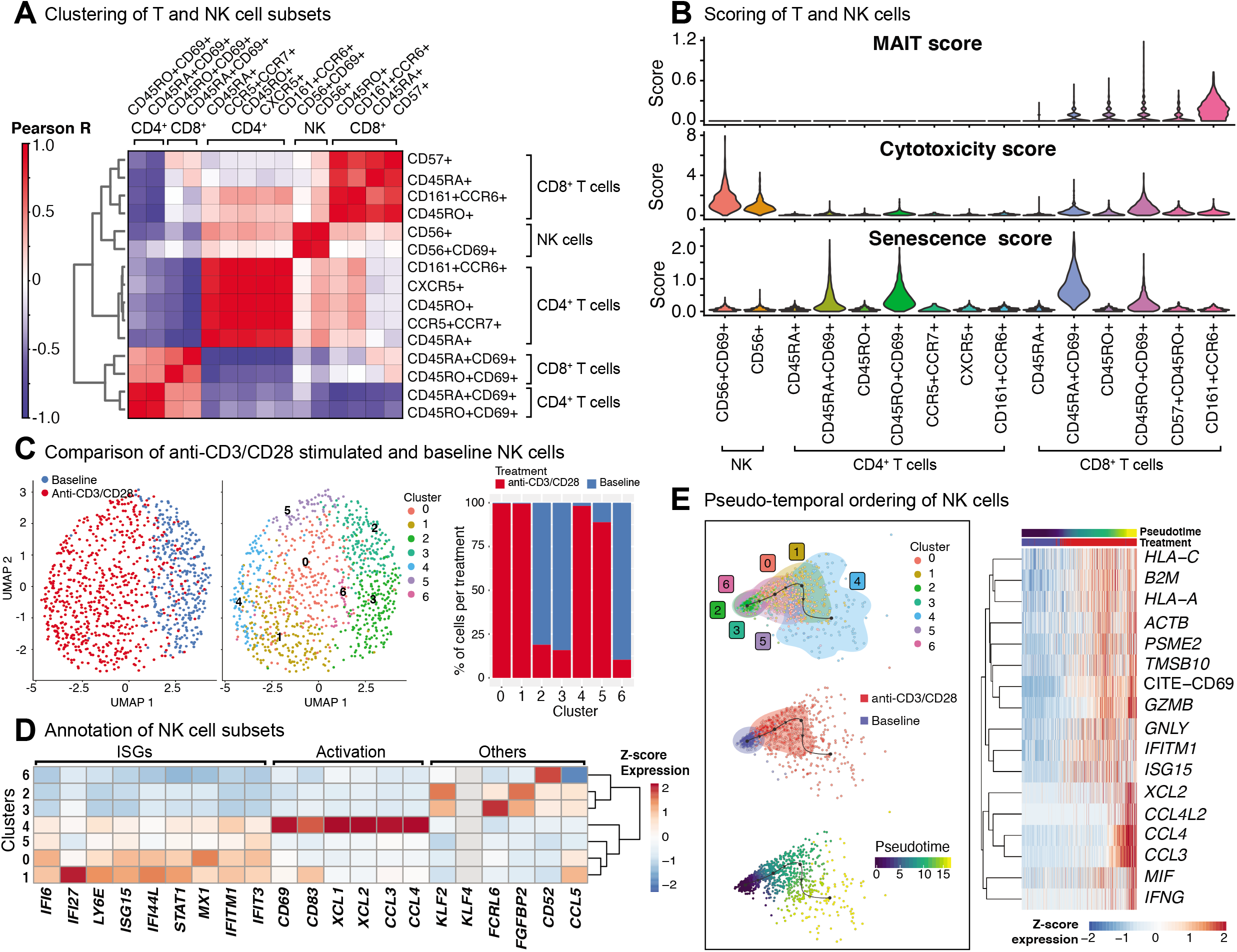
Indirect activation of NK cells. (A) Baseline and activated T cell and NK cell subsets were clustered together and the top 50 principal components were extracted. Pearson correlation values were then computed for each baseline and activated cell type and summarized in the heatmap. (B) Scoring of all T and NK cells was determined using transcripts associated with MAIT cell identity, cytotoxicity, and senescence (genes used are shown in Table S3). (C) Clustering of NK cells at baseline, and anti-CD3/CD28 conditions identifies 7 different subgroups (middle). In particular, anti-CD3/CD28-stimulated NK cells clustered distinctly from those at baseline (left). Distribution of NK cells in each cluster across three conditions (left). Note the enrichment of activated NK cells in clusters 0, 1, 4, and 5. (D) Heatmap of interferon stimulated genes (ISGs), activation-associated genes, and others used to annotate the NK sub-clusters. Heatmap values represent the average scaled expression of each gene for cells in the corresponding cluster. (E) (left) Pseudotime trajectory inference of NK cells at baseline and anti-CD3/CD28 conditions. (right) Genes and proteins whose expression patterns change with pseudotime in NK cells.

The significant effect of anti-CD3/CD28 treatment on the NK cell transcriptome (Fig. 4C; left) results from indirect activation, since neither TCR nor CD28 are expressed by NK cells. Unsupervised clustering of NK cells from baseline and anti-CD3/CD28 stimulation identified seven groups (Fig. 4C; middle), four of which were enriched in activated cells (~95% activated, Clusters 0, 1, 4, and 5) (Fig. 4C; right). Cells in these four clusters of activated NK cells had higher expression of interferon-stimulated genes (ISGs) including *LY6E, IFI6,* and *IFI27* (Fig. 4D) than NK cells at baseline. One of these clusters (cluster 4) highly expressed marker proteins for activation: *CD69* and *CD83* (35) (Fig. 4D). NK cells in cluster 4 expressed *XCL1* that facilitates their communication with DCs, as well as *CCL3* and *CCL4* (36) that enable them to recruit various immune cells including naive CD8^+^ T cells. In contrast, clusters 2, 3, and 6 cells were mostly from the baseline condition and enriched for *KLF2,* which encodes a transcription factor important for NK cell survival (37). Trajectory analyses of NK cells (baseline and activated) revealed that the cells at the end of the trajectory have the highest expression of interferon-gamma *(IFNG),* granzyme B *(GZMB),* CD69, and various chemokines *(CCL3, CCL4, XCL1, XCL2)* (Fig. 4E), resembling the profiles of cluster 4 cells. Expression of other interferon-inducible genes *(IFITM1, ISG15)* and granulysin *(GNLY)* peaked prior to the end of the trajectory, highlighting differences in timing and degree of NK subtype activation. Thus, scRNAseq of anti-CD3/CD28 activated PBMCs revealed that NK cells are indirectly activated with this condition that resulted in upregulation of cytotoxic molecules.

### Anti-CD3/CD28 activation of PBMCs results in B cell activation

Upon anti-CD3/CD28 treatment of PBMCs, B cells upregulated 383 genes and 4 cell surface proteins (CD69, CD25, ICOS, CD38) (Fig. 5A, Table S5). Increased transcription included ribosome and proteasome activity, oxidative phosphorylation machinery, and mitochondrial stress pathways. It also involved genes associated to humoral immune responses and Type 2 interferon signaling, such as *CD40, CXCL9, CXCL10,* and *IRF1* (Table S5). Resting and activated B cells clustered into six distinct groups (Fig. 5B; middle), where clusters 0 (comprising 25% of B cells) and 3 (comprising 15% of B cells) were specifically enriched in activated B cells (>90%) (Fig. 5B; right). Cells from the activated B cell clusters highly expressed ISGs *(IF44, IF44L, STAT1,* and *MX1*) and the activation marker *CD69* (Fig. 5C). Cluster 0 cells expressed genes associated with age-associated B cells (ABC) (38–40) and DN2 cells (41) (e.g., *TLR7, PRDM1, TGAX/*CD11c and *TBX2*1/T-bet). Distinct from ABCs and DN2 cells, they retained follicular (*CXCR5)* and conventional memory (*CD27)* markers. Cluster 3 cells expressed additional markers associated with DN2 cells, such as *FCRL5* and *TRAF5,* but lacked the expression of *TLR7, ITGAX,* and *TBX21.* Differences detected between these activated B cells and ABC/DN2 cells may reflect acute versus chronic activation of B cells as seen with aging and auto-immune diseases. Slingshot analyses revealed mostly linear activation trajectories for B cells, where the cells at the end of the trajectory had the highest expression of activation markers (CD25, CD69, and CD278) as well as genes encoding ribosomal proteins (Fig. 5D). Thus, scRNAseq of anti-CD3/CD28 activated PBMCs is providing new clues to the generation of DN2 cells, that are expanded in SLE (25,41).

**Fig. 5:**
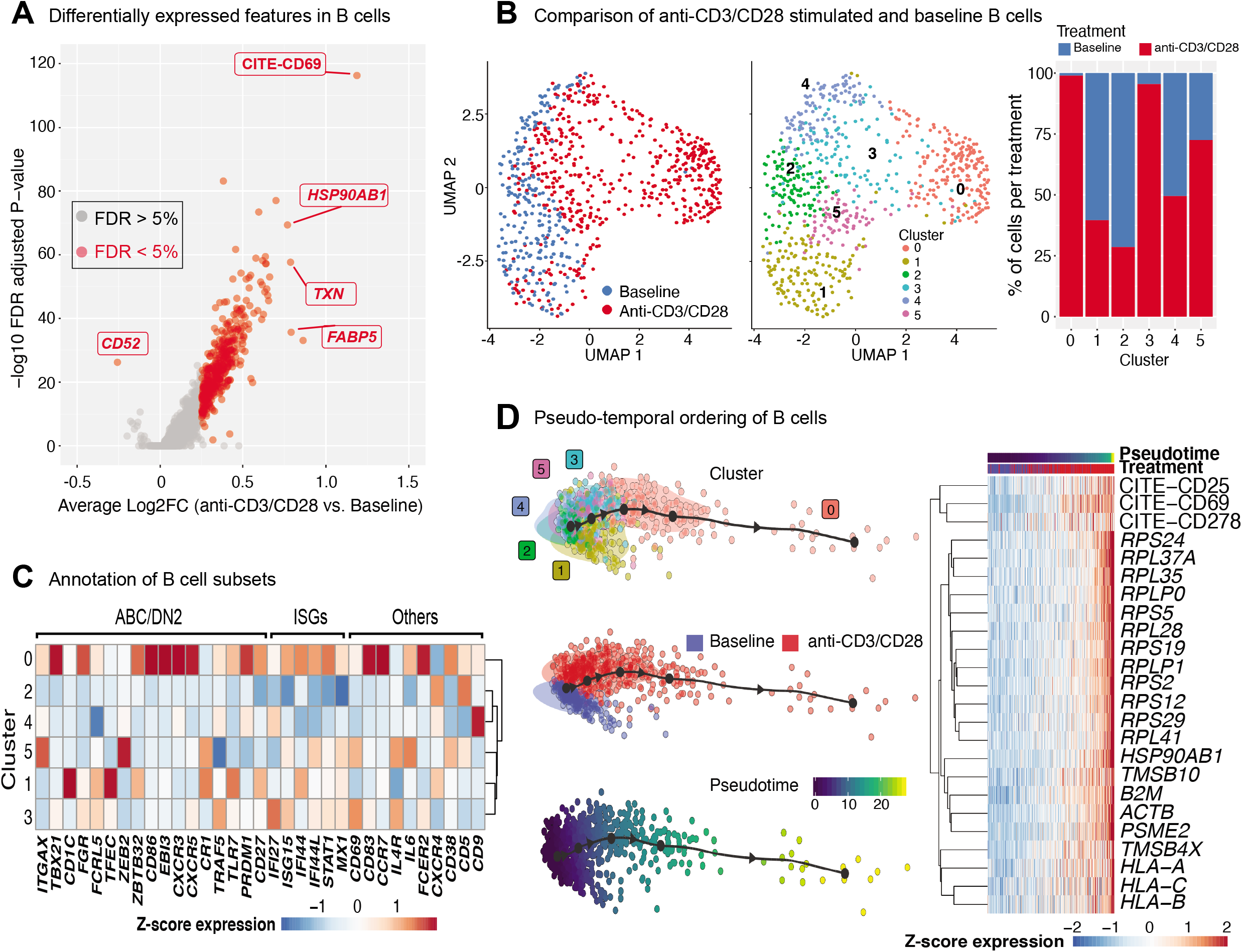
Indirect activation of B cells. (A) Volcano plot summarizing the genes induced in anti-CD3/CD28-activated B cells relative to those at baseline. (B) Analysis of B cells at baseline and after anti-CD3/CD28 stimulation revealed 6 subgroups (middle). Primarily, anti-CD3/CD28-stimulated B cells clustered separately from those at baseline (left). Distribution of B cells in each cluster across the two conditions (right). Activated B cells were mostly in clusters 0 and 3. (C) Heatmap of age-associated B cell (ABC)/double negative 2 (DN2), interferon stimulated genes (ISGs), and other genes used to annotate the distinct B cell subpopulations. Values in each heatmap represent average scaled gene expression of all cells in the corresponding cluster. (D) (left) Pseudotime trajectory inference of B cells at baseline and anti-CD3/CD28 conditions. (right) Genes and proteins with temporal patterns of expression in pseudo-temporal ordered B cells.

### LPS treatment of PBMCs only affects CD14^+^ monocytes and induces inflammation-associated genes

Baseline and LPS-activated PBMCs were jointly analyzed using the top 500 most variably expressed genes and proteins (Fig. 6A) and were annotated using the expression of marker proteins (Methods) (Fig. S3A). LPS stimulation did not lead to significant changes in PBMC cell compositions (Fig. 6B). Among PBMCs, only monocytes were activated by the LPS treatment (Fig. 6A, C) and showed upregulated expression of pro-inflammatory cytokine genes (e.g., *IL8, IL1B)* (Fig. S3B). To globally assess the enrichment of inflammation-associated transcripts in each cell type, we used a reference list of 249 inflammation-related genes from NanoString Technologies Inc. (Methods). Activated monocytes had significantly higher inflammation scores compared to baseline monocytes and all other cell types (Fig. 6D). Overall, 219 genes were differentially expressed in monocytes (125 up- and 94 down-regulated; FDR 5%) upon LPS treatment (Fig. 6E; Table S5). Functional enrichment analyses of these differentially expressed genes (DEGs) using gene ontology (GO) terms and KEGG pathways(42) revealed that LPS treatment activated pattern recognition receptor signaling pathways (Toll-like and NOD-like) and the pro-inflammatory NF-kappa B signaling pathway (FDR 5%; Table S6, Fig. S3C), including the up-regulation of important chemokines *(CCL3, CCL4)* and cytokines *(IL1B, IL6)* (Fig. 6F). In contrast, LPS stimulation decreased the expression of genes associated with antigen processing and presentation *via* MHC class II and phagosome activity (Fig. 6F, Table S6).

**Fig. 6:**
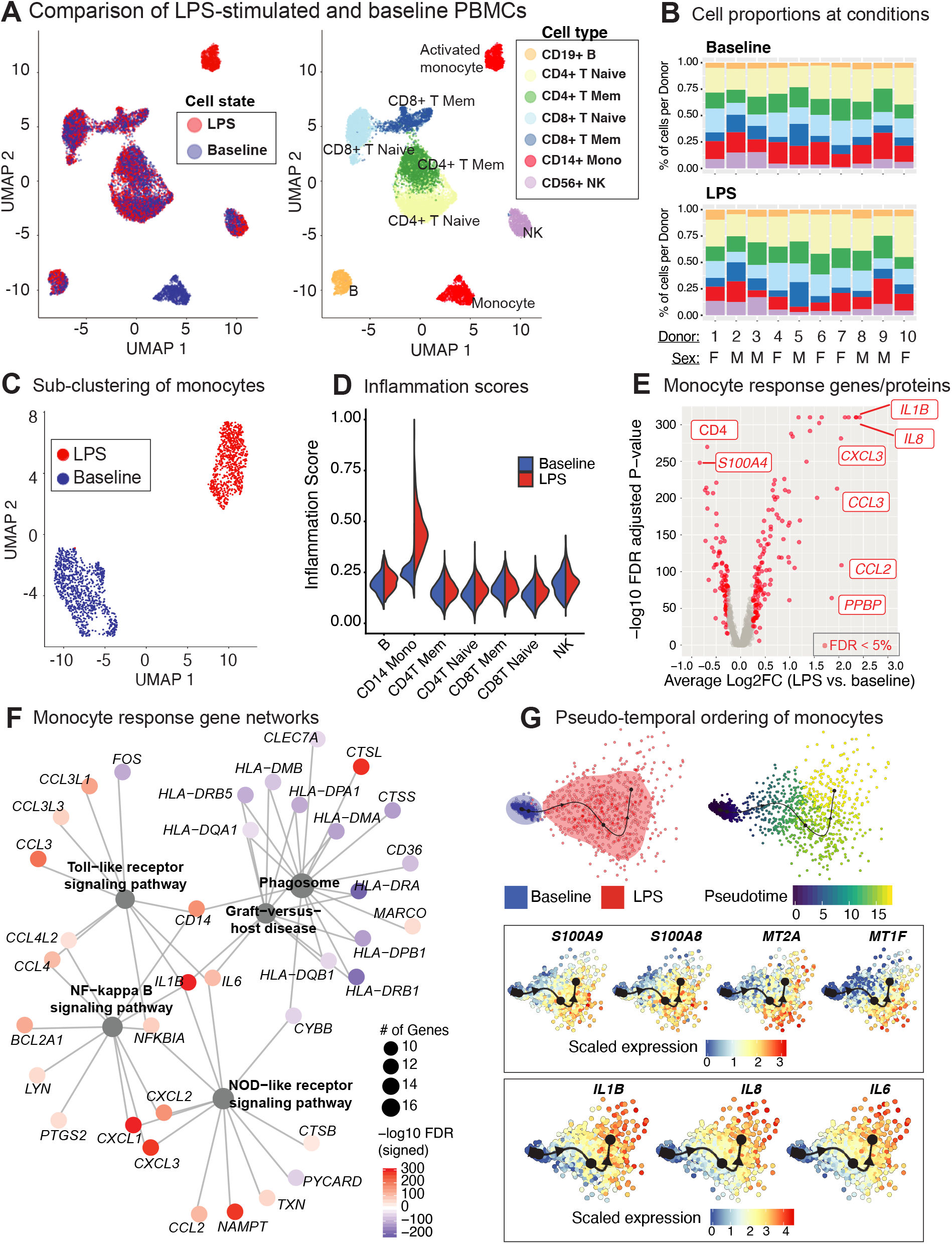
LPS stimulation specifically activates monocytes and induces pro-inflammatory responses. (A) Clustering of PBMCs under baseline and upon LPS stimulation (left). Cell types were annotated using the expression of cell surface proteins (right). (B) Cell type compositions at baseline (top) and LPS condition (bottom) across 10 donors. Refer to panel A for cell types’ color coding. (C) Clustering of CD14^+^ monocytes separates baseline and LPS-stimulated cells. (D) Inflammation scores of all cells under two conditions. LPS-activated monocytes have the highest expression levels of inflammation-associated transcripts. (E) Differential expression analysis of LPS-stimulated and baseline monocytes revealed 125 induced and 94 reduced genes respectively (FDR 5%), as well as 3 induced and 3 reduced proteins, respectively. (F) Enriched KEGG pathways (FDR 5%) for LPS response genes. Node size for each pathway represents the number of genes in the pathway that were also induced in LPS-stimulated monocytes. Node color for genes indicate induction (red) or reduction (blue) upon LPS treatment. LPS-activated monocytes upregulate genes associated with pattern recognition receptor signaling pathways (Toll-like receptor, NOD-like receptor) and inflammation (NF-kB signaling), but also downregulate genes associated with antigen presentation and phagosome activity. (G) Trajectory (pseudotime) inference of monocytes under baseline and LPS conditions identified genes with heterogeneous activity in stimulated monocytes. Cells are color-coded with respect to activation status (top left) and inferred pseudotimes (top right). Expression levels of selected genes were overlaid on cells sorted in the activation trajectory (bottom).

### LPS-activated monocytes exhibit two distinct transcriptional states

The Slingshot (34) algorithm revealed that the trajectory of monocytes followed a linear transition from resting to activated states (Fig. 6G; top, Fig. S3D). LPS-activated monocytes showed increased variation in their transcriptional responses and were positioned at the end of the trajectory (Fig. 6G; top). Genes that are important for defining monocyte trajectories included inflammatory cytokines *(IL6, IL8, IL1B),* metallothionein genes involved in zinc transport *(MT1F* (43)) and oxidative stress *(MT2A* (44)), as well as alarmins *(S100A8/S100A9),* which induce pro-inflammatory cytokine production (45) (Fig. S3E). Alarmins and metallothionein gene expression peaked in early stages (pseudotime values 5 up to 15), whereas pro-inflammatory cytokines (e.g., *IL1B, IL6, IL8)* peaked in late stages (pseudotime >=15) (Fig. 6G; bottom, Fig. S3E). These LPS-induced cellular states and changes were observed consistently across donors (Fig. S3F). To further dissect this heterogeneity, we clustered baseline and activated monocytes using k-means clustering (k=4). Baseline monocytes clustered separately from LPS-treated ones (Fig. 7A, S4A), which were split into 3 clusters referred to as LPS1, LPS2, and LPS3 states. These clusters coincided with the pseudotime values (Fig. S4B), where LPS1 and LPS2 states mapped to the early activation state and LPS3 mapped to the later inflammatory state (Fig. S4B). These four monocyte clusters were detected in all individuals (Fig. 7B); inflammation scores of clusters (Methods) increased in agreement with their position in the trajectory, LPS3 having the highest inflammation score (Fig. 7C). LPS2 and LPS3 states had the most distinct transcriptional response profiles, with 50 and 47 genes uniquely induced in these states, respectively, compared to baseline (Fig. S4C, S4D, Table S7).

**Fig. 7:**
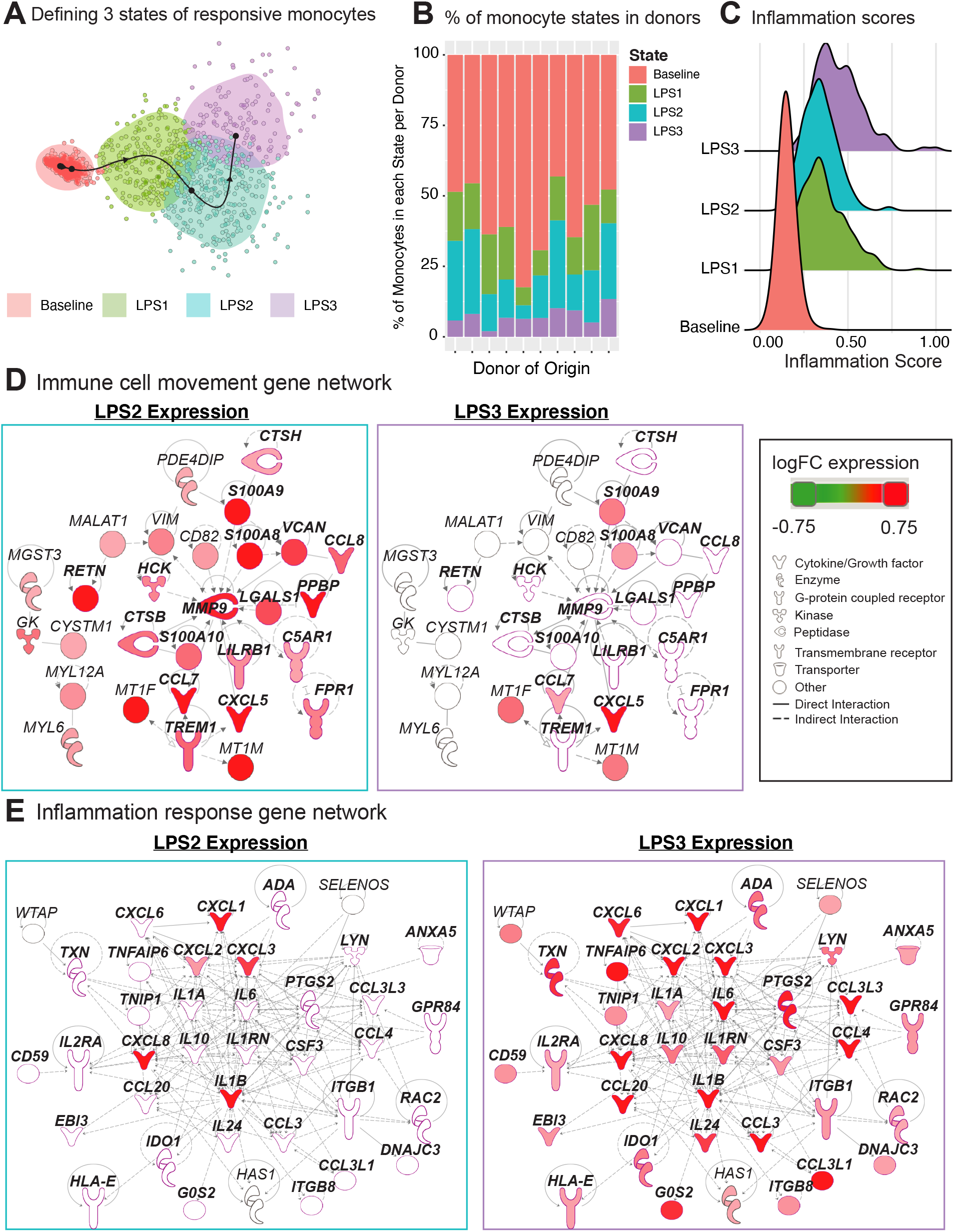
Pseudo-temporal ordering of monocytes reveals distinct LPS-activation states. (A) K-means clustering (n = 4) of the monocyte trajectory divides baseline cells into one group and activated cells into three groups (LPS1, LPS2, LPS3). (B) Proportions of monocytes found in each of the four states across the 10 donors. (C) Inflammation scores of monocytes in distinct states. LPS3 state monocytes show the highest average inflammation score. (D) Network of genes induced in activated monocytes from LPS2 state using Ingenuity Pathway Analysis (IPA). Molecules in bold are associated with immune cell migration and chemotaxis. (E) Network of genes induced in LPS3 activated monocytes. Bolded molecules are associated with inflammatory responses. For both networks, molecules with different functions were depicted with different shapes, as provided by IPA. Molecules are colored based on their log2 fold change in expression in that state vs. all other baseline and activated states. Direct and indirect gene interactions are depicted by solid and dashed lines respectively.

Network analyses of genes induced in LPS2 and LPS3 states using Ingenuity Pathway Analysis (IPA; Methods) revealed that LPS2 genes included alarmins *(S100A8, S100A9)* and genes associated with cell movement and trafficking (Fig. 7D; genes in bold text, pink outline), such as various chemoattractants *CXCL7 (PPBP), CCL7,* and *CCL8.* In contrast, LPS3 genes included pro-inflammatory interleukins (e.g., *IL1A, IL1B, IL6),* macrophage inflammatory protein-1-alpha (*CCL3*), - 1-beta (*CCL4*), and −3 (*CCL20*), and tumor necrosis-associated factors *(TNFAIP6, TNIP1)* (Fig. 7E). Interestingly, in LPS3, we also observed the induction of anti-inflammatory genes including *IL10, HLA-E* - an MHC class I molecule with inhibitory effects on cytokine production in monocytes (46), *IDO1* - an immunosuppressive enzyme - and one of its regulators *PTGS2* (47), suggesting a feedback loop to control increased inflammation. Pathway enrichment analysis using IPA disease and function terms verified that pro-inflammatory signals emerge at LPS2 state and peak at the later LPS3 state (Fig. S4E). At baseline, monocytes showed higher relative expression of ribosomal protein genes (Fig. S4F) and genes involved in mRNA processing, translation, and protein synthesis. Thus, scRNAseq analysis further delineates how LPS activates monocytes into early and late inflammatory states.

## Discussion

*In vitro* activation of circulating immune cells have been widely used for immune monitoring and identification of disease mechanisms since 1980s (8,48). However, such assays are generally limited in scope and often only provide information regarding a single cell type. A holistic study of circulating immune cell responses to activators will uncover how the system as a whole respond to an immune challenge and how cell-cell interactions contribute to these responses. Hence these studies will bring us closer to a better understanding of immune cell responses in health and disease. CITE-seq enables precise studies of in-vivo and in-vitro immune cell responses. However, these assays are costly to generate and harbor technical challenges that might confound data interpretation (i.e., multiplets, batch effects). Herein we describe a cost-effective and in-depth study to uncover responses of peripheral blood mononuclear cells (PBMCs) from 10 healthy donors (5 men, 5 women) to two widely used activators - anti-CD3/CD28 and LPS - to activate adaptive and innate cells, respectively. PBMCs were profiled using CITE-seq technology with 39 antibodies before and after activation; libraries were multiplexed using both cell hashing and genotypes, which reduced costs and batch effects. This design enabled studying activated cells in a cost-effective manner while also detecting and labeling multiplets from i) different conditions, ii) different individuals; and iii) different cell types. We observed an increase in biological multiplets upon activation, potentially due to increased cell-cell interactions, which should be taken into consideration in future study designs to decouple biological (i.e., cellular interactions) from technical (i.e., due to superloading) multiplets (49,50). One caveat of multiplexing many libraries is the limited number of cells detected per donor (~1600 cells), which was enough to study responses of major cell populations (e.g., CD14^+^ monocytes), but not that of less abundant ones (e.g., CD16^+^ monocytes, dendritic cells). Multiplexing fewer samples can effectively address this caveat in future studies.

Anti-CD3/CD28 treatment activated all classes of lymphocytes either directly (T cells) or indirectly (B, NK cells). Upon activation, co-stimulatory molecules (CD28, CD278) and activation markers (CD69, CD25) were expressed on the surface of both naïve and memory T cells. Transcriptomic changes in naïve T cells were the greatest likely due the fact that these cells start from a ‘lower transcriptional state’ compared to their memory counterparts as they need to differentiate and initiate clonal expansion (51). Activated T cells highly expressed senescence (e.g., *CDKN1A* - encoding p21, *CDKN2A* - encoding p16, *SESN2,* and *PRKAA1),* effector and cytotoxic molecules (e.g., *GZMA/B/H, IFNG).* Using CITE-seq antibodies, under baseline conditions, we detected distinct subsets of memory cells (e.g., Th subsets, MAIT). However, these subsets were not detected upon activation, potentially due to the internalization of cell surface molecules and increased transcriptional similarity between memory subsets.

Indirect activation of NK cells *via* anti-CD3/CD28 activated 74 genes, most of which were shared with T cell responses including CD69, interferon stimulated/response, and cytotoxic genes. Scoring of single cells using different gene sets showed that activated NK cells had the highest scores for cytotoxicity whereas activated T cells had highest scores for senescence. Clustering and trajectory analyses revealed heterogeneity among activated NK and B cells. Late stage activated NK cells resembled previously described ‘helper’ NK cells and expressed CD83 protein (35), whereas late stage activated B cells expressed molecules that have been expressed in B cells associated with aging and auto-immune diseases (*TLR7, PRDM1, ITGAX/CD11c, TBX21/T-bet)* (39–41). However, compared to DN2s and ABCs, activated B cells retained follicular and conventional memory markers (*CXCR5* and *CD27*), likely as a reflection of acute versus chronic activation. Future single cell activation studies from different populations should help further characterize these B cells.

LPS stimulation specifically activated monocytes and pseudo-temporal analyses uncovered two states of activated monocytes: i) a first state that includes pro-inflammatory alarmins *(S100A8/A9)* and molecules to attract and interact with other immune cells *(PPBP, CCL7, CCL8);* and ii) a second, possibly more terminated inflammatory state, with increased expression of *bona fide* inflammatory cytokines *(IL1B, IL6, IL8,* and *IL24)* and potential anti-inflammatory molecules *(IL-10, SOCS3, HLA-E,* and *IDO1)* to mitigate the effects of increased inflammation. A similar phenomenon was noted in human blood dendritic cell subsets in which cellular responses to influenza virus infection resulted in three waves of chemokine secretion to attract effector immune cells (52). Induction of interferon response genes and TFs (STAT1 and IRF1) previously reported in IFN-g-primed (24 hr) (53) and LPS-induced (0-4 hrs) monocytes (54), was not observed in our study, potentially due to the 24 hour stimulation, which is a later time-point in the activation cascade. Anti-CD3/CD28 treatment reduced the numbers of monocytes in PBMCs likely due to the induced apoptosis *via* activated T cells (28).

In summary, this study provides an experimental and computational framework to carefully study activated immune cells using single cell methods by multiplexing to handle batch effects and cut costs and by quantifying protein expression to identify cell types, cell states, and multiplet cells. The data, analyses, and application developed here will be important resources for future studies such as exposure of immune cells to antigens, microbes, adjuvants and other targeting agents. It will guide the design of subsequent experiments to further expand our understanding of cellular responses to additional *in vivo* and *in vitro* activation conditions in healthy and diseased individuals. Overall this study provides a foundation for future immune activation experiments using single cell technologies. The data is freely shared via Human Cell Atlas platform as well as via an interactive R shiny application (https://czi-pbmc-cite-seq.jax.org/).

## Methods

### Blood collection

Blood was collected at the UConn Health Clinical Research Center (CRC) at 263 Farmington Avenue Farmington, CT 06030-3805 following informed consent of participants for a UConn Health IRB-approved protocol (IRB Number: 18-151J-2). Approximately 50ml of blood was collected from each donor. Samples were collected BD vacutainer tubes containing Acid Citric Dextrose (ACD) Solution A, and peripheral blood mononuclear cells (PBMC) were isolated immediately after.

### Isolation and stimulation of PBMCs

Peripheral blood mononuclear cells (PBMCs) were isolated with gradient centrifugation using Lymphoprep (StemCell Technologies) and the red blood cells were removed by inclubating the cells with RBC lysis buffer (Tonbo Biosciences). Atfer that, the PBMCs were immediately cultured using activation conditions described below for 24 hours at 1 million cells/ml density in complete RPMI (RPMI supplemented with 10% heat-inactivated Fetal bovine serum, 100U/ml Penicillin/Streptomycin, 2mM L-glutamine, 10mM HEPES, 0.1mM Non-essential amino acids, 1mM Sodium pyruvate) at 37°C + 5% CO2. The PBMCs were cultured in three different ways: 1) Plate-bound anti-CD3 + anti-CD28 (clones OKT3 and CD28.2 at 2μg/ml each - Tonbo Biosciences) - plates were coated with Goat anti-mouse IgG (clone Poly4053 at 10μg/ml - BioLegend) for 1 hour at 37°C, washed, then coated with both anti-CD3 and anti-CD28 also for 1 hour at 37°C. After that cells were added to the plates for stimulation; 2) lipopolysaccharide (LPS) (10ng/ml - InvivoGen); and 3) Complete medium only as control. After 24 hours in culture, an aliquot of each sample was stained for flow cytometry analysis and the remaining cells were harvested and frozen for subsequent CITE-seq analysis and genotyping.

### Flow cytometry and antibodies

PBMCs cells were incubated with Ghost Dye™ Violet 510 (Tonbo Biosciences) to stain dead cells for 10 min at room temperature (RT). After, Fc receptors were blocked for 5 min at RT using FcR-blocking reagent (Miltenyi Biotec) and then incubated with fluorochrome-conjugated surface antibodies for 15 min at RT. The surface antibodies CD3 (UCHT1), CD11b (M1/70), CD14 (HCD14), CD16 (3G8), CD19 (HIB19), CD20 (2H7), CD25 (BC96), CD27 (O323), CD69 (FN50), HLA-DR/MHC class II (L243) were all from BioLegend and CD80 (2D10.4) from eBioscience. FACS analysis was performed on a BD LSR Fortessa (BD Biosciences) and data were analyzed with FlowJo software (Version 10.2, TreeStar).

### CITE-seq data generation with Cell Hashing

CITE-seq data was generated from 30 samples (10 individuals, 3 experimental conditions) by multiplexing to reduce batch effects and costs using cell hashing (55) (for experimental conditions) and genotype-based demultiplexing (27) (for individual donors). For CITE-seq experiments, antibody-oligo conjugates against 39 surface proteins (e.g., CD8, CD45RA, CD4, CD3, CCR7, CD69) as well as the Cell Hashing Antibodies (HTOs) were purchased from BioLegend. PBMCs from different donors were independently stained with one of our HTO-conjugated antibody pools (based on the experimental condition) and a pool of 39 immunophenotypic markers for CITE-seq. Equal numbers of cells from all 30 samples were pooled and super-loaded onto 10 lanes of 10x Genomics Single Cell 3P v2 (10x Genomics, USA), yielding approximately 25K cells per lane. All libraries were sequenced together on Illumina NovaSeq S4 flowcells.

### Genotyping of PBMCs

PBMCs from 10 donors were genotyped using Infinium Omni2.5-8 v1.3 BeadChip Array (Illumina) according to manufacturer’s instructions. We mapped the Illumina array probe sequences to hg19 genome assembly and excluded potential problematic ones as in (56,57). Furthermore, we filtered out SNPs with an alternate allele frequency difference with 1000G EUR samples > 20%, or palindromic SNPs with a minor allele frequency > 20%, genotype missingness > 2.5%, Hardy-Weinberg p-value < 10-4. Finally, 1,147,545 genotyped SNPs were considered for further analyses.

### Sample demultiplexing using Cell Hashing

Three hashtag oligonucleotides (HTO) were used to multiplex samples from different experimental conditions (e.g., HTO A for baseline samples; HTO B for Anti-CD3/CD28 treated samples). Cell HTOs were counted in raw sequencing data using CITE-seq-Count version 1.4.2. (55). CITE-seq-Count was used to count HTOs for each cell using recommended parameter settings (“-cells 25000 -cbf 1 -cbl 16 -umif 17 -umil 26 --max-error 3”). HTO counts per cell were normalized with respect to the total read counts for that cell and scaled and log2 transformed. Distributions of read counts for each HTO across all cells were bimodal. To classify whether a cell has detectable counts for an HTO, we fit a Gaussian mixture model (with 2 mixture components) for each HTO using mclust R package (58) (version 5.4.5). Based on these models, we labeled each cell as: i) empty: if there are no sufficient read counts for any HTO; ii) singlet: only if counts for a single HTO were detected; iii) multiplets: if there are detectable counts for 2 (“doublet”) or 3 (“triplet”).

### Sample demultiplexing with Demuxlet

Genotypes were used to demultiplex the donor of origin for each sample using Demuxlet version 2 (27) with default parameters. For this we used 167,703 genotyped SNPs overlapping exonic regions in the genome. Cells annotated to one individual were retained, while “ambiguous” and “doublet” cells were discarded from further analyses.

### RNA quantification

Single cell RNA-sequencing libraries were prepared using the Chromium Single Cell 3’ Reagent Kit (v2 Chemistry) according to the user guide (10X Genomics). Resulting libraries were sequenced using Illumina NovaSeq S4 flowcells to an average depth of ~150,000 reads per cell. Reads were aligned to human reference sequence GRCh37/hg19 and unique molecular identifiers (UMIs) were quantified using the Cell Ranger “count” software (10X Genomics, version 2.1.0) and with additional parameters “--expect-cells=25000”. Cell barcodes with fewer than 400 detected genes, and those not identified to be singlets via cell hashing or Demuxlet were discarded from analyses.

### Protein quantification

Read counts for antibody derived tags (ADTs) for 39 distinct antibodies were measured from raw sequencing reads using CITE-seq-Count with the following parameters: “-cells 25000 -cbf 1 -cbl 16 -umif 17 -umil 26 --max-error 3”. Raw counts were normalized using a centered log ratio approach using Seurat R package version 3.0.2 (59).

### Multiplet cell removal using surface protein expression

Number of multiplet cells increases upon activation, due to the increase in cell-cell interactions. To remove such multiplets, we took advantage of cell surface protein expression levels and identified cells that co-expressed multiple cell-type-specific proteins. First, we determined which cells were “positive” and “negative” for each protein using the HTODemux function from Seurat (59). Since some immune cells co-express certain cell surface markers (e.g., NK and CD8^+^ T cells both express CD8), we devised a multiplet removal protocol for each cell type. For B cell multiplets, we filtered out cells that express CD19 in addition to one of the following i) a T cell marker (CD3, CD4, CD8); ii) an innate cell marker (CD14, CD16, CD56). For CD14^+^ monocyte multiplets, we removed cells that express CD14 and i) a T cell marker (CD3, CD8, CD28); ii) NK marker CD56; iii) a B cell marker (CD19, CD20). For NK cell multiplets, we removed cells that are positive for CD56 and one or more of the following i) T cell markers (CD3, CD4, CD28); ii) monocyte markers (CD14, CD11c); iii) B cell markers (CD19, CD20). For CD4^+^ T cell multiplets, we removed cells that express CD4 markers (CD3, CD4) and one or more of the following i) CD8 (for CD8^+^ T), CD56 (for NK), CD19, CD20 (for B cells), CD14, CD16 (for monocytes/NK). Lastly, for CD8^+^ T cell multiplets, we removed cells that co-expressed CD8 markers (CD3, CD8) and one or more of the following i) CD4 (for CD4^+^ T cells), ii) CD56 (for NK), iii) CD19, CD20 (for B cells), and iv) CD14, CD16 (for monocytes/NK). After all cells are removed we obtained comparable number of single cells for each individual upon 3 conditions. These cell numbers correlated significantly with cell numbers inferred from flow cytometry data (R=0.97).

### Cell clustering

Cells were clustered using top 500 most variable genes and proteins, which are determined by fitting a line to the relationship of the variance and mean using a local polynomial regression (loess). Protein/gene expression values were standardized using the observed normalized mean and expected variance (given by the fitted line). Feature variance was then calculated on the standardized values after clipping to a maximum of the square root of the number of cells. After scaling and centering the feature values, principal component analysis (PCA) was performed on the top 500 variable features. Following this, uniform manifold approximation and projection (UMAP) was applied on the top 20 principal components to reduce the data to two dimensions. Next, a shared nearest neighbor (SNN) graph was created by calculating the neighborhood (Jaccard index) overlap between each cell and its 20 (default) nearest neighbors. To identify cell clusters, we used a SNN modularity optimization based clustering algorithm (60).

### Differential expression

To identify cell-type-specific response genes and proteins, we conducted pairwise Wilcoxon Rank Sum tests between each PBMC cell-type under stimulation and baseline conditions (e.g., LPS stimulated monocytes vs. baseline monocytes). In each differential expression analysis, genes detected in at least 10% of each cell group were considered. Features with an absolute difference in expression > log2(0.25) and false discovery rate < 5% were regarded as significant results.

### Single cell pseudo-temporal trajectory inference

Benchmarking and inference of single cell pseudotime trajectories were performed with the R package *dyno* (version 0.1.1) (61). Briefly, for each cell-specific response (e.g., monocytes induced by LPS) trajectories were inferred using Slingshot (the method determined to minimize error and maximize scalability and accuracy). Pseudo-temporal ordering of cells were derived using the corresponding baseline and activated cells as well as the genes and ADTs previously determined to be differentially induced (FDR < 5%) between the two groups of cells.

### Ingenuity Pathway Analysis

Response genes were uploaded to Qiagen’s Ingenuity Pathway Analysis (IPA) version 01-16 system to build gene networks for each gene set. Networks were constructed using the Connect tool with “Human” as the species parameter and otherwise default parameters to map direct and indirect interactions between molecules as described in the Ingenuity Knowledge Base (IKB). IPA’s core analysis function was also used to annotate each response gene set and resultant networks with canonical pathways, relevant diseases, and biological functions.

### Single cell scoring

Genes associated with inflammation were obtained from NanoString Technologies, Inc. (cite, https://www.nanostring.com/products/gene-expression-panels/gene-expression-panels-overview/ncounter-inflammation-panels) and used to calculate an inflammation score for PBMCs at baseline and LPS-stimulated conditions. Feature counts for each cell were divided by the total counts for that cell, multiplied by a scale factor of 10,000, and then log normalized. Normalized read counts for all inflammation-associated genes were divided by the normalized counts of all transcripts and then min-max scaled between 0 and 1. Gene lists (Table S3) were used to score cytotoxicity, MAIT or senescence expressions across T and NK cell clusters. To do so, we calculated the mean expression for each cell, within each cluster.

### Pathway analysis of stimulation-response genes

Differentially expressed genes for each cell-type-response to stimulation were functionally annotated using the R package *clusterProfiler* (version 3.14.0) (42). The Kyoto Encyclopedia of Genes and Genomes (KEGG), gene ontology (GO), and WikiPathways databases were used to determine association with particular biological processes, diseases, and molecular functions. The top pathways with an FDR-adjusted p-value <5% were summarized in the results.

## Author Contributions

D.U., A.W., M.S., P.S., designed the experimental study design. M.S, P.S., A.W., and J.G. processed PBMC samples and the sequencing of libraries. N.L. and D.N.-B., performed bioinformatics analyses of the data. N.L., D.U, A.W, D.N.-B. M.L.S., V.P., and J.B. interpreted the data. N.L, J.B., A.W., and D.U. wrote the manuscript. N.L., J.B., D.N.-B., A.W. and D.U. designed the corresponding figures. N.L. created the R shiny web application. All authors edited and reviewed the manuscript prior to submission.

## Acknowledgements

We thank all members of the Stitzel and Ucar labs for constructive feedback and discussion on this study and manuscript. We also thank members of the Human Cell Atlas data wrangler team for assistance with depositing the data associated with this manuscript.

## Supplementary Material

Table S1: Barcoding information for the 39 antibodies (CITE-seq panel) and cell hashing oligonucleotides (HTOs) used to multiplex treatment information.

Table S2: Sequencing quality control information for RNA-seq, cell hashing, and protein quantification data.

Table S3: Reference gene lists used to compute single cell scores for inflammation, NK/cytotoxicity, MAIT, TEMRA, and senescence.

Table S4: Lists of anti-CD3/CD28 response genes/proteins that are shared or unique to T and NK cell subsets.

Table S5: Differential gene and protein expression analysis results for each cell-type at the specified condition vs. that at baseline condition.

Table S6: Pathway enrichment analyses for monocyte responses to LPS.

Table S7: Genes induced in each of the baseline and LPS-responsive monocyte states (LPS1, LPS2, and LPS3).

## Data Availability

The accession numbers for human PBMC CITE-seq data reported in this paper are: European Variation Archive project PRJEB40448 and analyses ERZ1631464. The R shiny web application to visualize and interact with the data generated in this study is freely available at https://czi-pbmc-cite-seq.jax.org/.

## References

1. Sen P, Kemppainen E, Orešič M. Perspectives on Systems Modeling of Human Peripheral Blood Mononuclear Cells. Front Mol Biosci (2018) 4: doi:10.3389/fmolb.2017.00096

2. Schmiedel BJ, Singh D, Madrigal A, Valdovino-Gonzalez AG, White BM, Zapardiel-Gonzalo J, Ha B, Altay G, Greenbaum JA, McVicker G, et al. Impact of Genetic Polymorphisms on Human Immune Cell Gene Expression. Cell (2018) 175:1701–1715.e16. doi:10.1016/j.cell.2018.10.022

3. Kotliarov Y, Sparks R, Martins AJ, Mulè MP, Lu Y, Goswami M, Kardava L, Banchereau R, Pascual V, Biancotto A, et al. Broad immune activation underlies shared set point signatures for vaccine responsiveness in healthy individuals and disease activity in patients with lupus. Nature Medicine (2020) 26:618–629. doi:10.1038/s41591-020-0769-8

4. Moskowitz DM, Zhang DW, Hu B, Le Saux S, Yanes RE, Ye Z, Buenrostro JD, Weyand CM, Greenleaf WJ, Goronzy JJ. Epigenomics of human CD8 T cell differentiation and aging. Sci Immunol (2017) 2: doi:10.1126/sciimmunol.aag0192

5. Banchereau R, Hong S, Cantarel B, Baldwin N, Baisch J, Edens M, Cepika A-M, Acs P, Turner J, Anguiano E, et al. Personalized Immunomonitoring Uncovers Molecular Networks that Stratify Lupus Patients. Cell (2016) 165:551–565. doi:10.1016/j.cell.2016.03.008

6. Persson U, Johansson SG. Concanavalin A-induced activation of human T cell subpopulations. Int Arch Allergy Appl Immunol (1984) 75:337–344. doi:10.1159/000233644

7. Trickett A, Kwan YL. T cell stimulation and expansion using anti-CD3/CD28 beads. Journal of Immunological Methods (2003) 275:251–255. doi:10.1016/S0022-1759(03)00010-3

8. Smith-Garvin JE, Koretzky GA, Jordan MS. T Cell Activation. Annual Review of Immunology (2009) 27:591–619. doi:10.1146/annurev.immunol.021908.132706

9. Rossol M, Heine H, Meusch U, Quandt D, Klein C, Sweet MJ, Hauschildt S. LPS-induced cytokine production in human monocytes and macrophages. Crit Rev Immunol (2011) 31:379–446. doi:10.1615/critrevimmunol.v31.i5.20

10. Cepika A-M, Banchereau R, Segura E, Ohouo M, Cantarel B, Goller K, Cantrell V, Ruchaud E, Gatewood E, Nguyen P, et al. A multidimensional blood stimulation assay reveals immune alterations underlying systemic juvenile idiopathic arthritis. J Exp Med (2017) 214:3449–3466. doi:10.1084/jem.20170412

11. Piasecka B, Duffy D, Urrutia A, Quach H, Patin E, Posseme C, Bergstedt J, Charbit B, Rouilly V, MacPherson CR, et al. Distinctive roles of age, sex, and genetics in shaping transcriptional variation of human immune responses to microbial challenges. PNAS (2018) 115:E488–E497. doi:10.1073/pnas.1714765115

12. Thomas S, Rouilly V, Patin E, Alanio C, Dubois A, Delval C, Marquier L-G, Fauchoux N, Sayegrih S, Vray M, et al. The Milieu Intérieur study - an integrative approach for study of human immunological variance. Clin Immunol (2015) 157:277–293. doi:10.1016/j.clim.2014.12.004

13. Molony RD, Nguyen JT, Kong Y, Montgomery RR, Shaw AC, Iwasaki A. Aging impairs both primary and secondary RIG-I signaling for interferon induction in human monocytes. Sci Signal (2017) 10: doi:10.1126/scisignal.aan2392

14. Shaw AC, Goldstein DR, Montgomery RR. Age-dependent dysregulation of innate immunity. Nat Rev Immunol (2013) 13:875–887. doi:10.1038/nri3547

15. Shaw AC, Panda A, Joshi SR, Qian F, Allore HG, Montgomery RR. Dysregulation of human Toll-like receptor function in aging. Ageing Research Reviews (2011) 10:346–353. doi:10.1016/j.arr.2010.10.007

16. Qian F, Guo X, Wang X, Yuan X, Chen S, Malawista SE, Bockenstedt LK, Allore HG, Montgomery RR. Reduced bioenergetics and toll-like receptor 1 function in human polymorphonuclear leukocytes in aging. Aging (Albany NY) (2014) 6:131–139.

17. Gowthaman U, Chen JS, Zhang B, Flynn WF, Lu Y, Song W, Joseph J, Gertie JA, Xu L, Collet MA, et al. Identification of a T follicular helper cell subset that drives anaphylactic IgE. Science (2019) 365: doi:10.1126/science.aaw6433

18. Papalexi E, Satija R. Single-cell RNA sequencing to explore immune cell heterogeneity. Nature Reviews Immunology (2018) 18:35–45. doi:10.1038/nri.2017.76

19. Regev A, Teichmann SA, Lander ES, Amit I, Benoist C, Birney E, Bodenmiller B, Campbell P, Carninci P, Clatworthy M, et al. The Human Cell Atlas. Elife (2017) 6: doi:10.7554/eLife.27041

20. Pereira BI, De Maeyer RPH, Covre LP, Nehar-Belaid D, Lanna A, Ward S, Marches R, Chambers ES, Gomes DCO, Riddell NE, et al. Sestrins induce natural killer function in senescent-like CD8+ T cells. Nat Immunol (2020) 21:684–694. doi:10.1038/s41590-020-0643-3

21. Szabo PA, Levitin HM, Miron M, Snyder ME, Senda T, Yuan J, Cheng YL, Bush EC, Dogra P, Thapa P, et al. Single-cell transcriptomics of human T cells reveals tissue and activation signatures in health and disease. Nature Communications (2019) 10: doi:10.1038/s41467-019-12464-3

22. Shalek AK, Satija R, Shuga J, Trombetta JJ, Gennert D, Lu D, Chen P, Gertner RS, Gaublomme JT, Yosef N, et al. Single-cell RNA-seq reveals dynamic paracrine control of cellular variation. Nature (2014) 510:363–369. doi:10.1038/nature13437

23. Villani A-C, Satija R, Reynolds G, Sarkizova S, Shekhar K, Fletcher J, Griesbeck M, Butler A, Zheng S, Lazo S, et al. Single-cell RNA-seq reveals new types of human blood dendritic cells, monocytes, and progenitors. Science (2017) 356: doi:10.1126/science.aah4573

24. Horns F, Dekker CL, Quake SR. Memory B Cell Activation, Broad Anti-influenza Antibodies, and Bystander Activation Revealed by Single-Cell Transcriptomics. Cell Reports (2020) 30:905–913.e6. doi:10.1016/j.celrep.2019.12.063

25. Nehar-Belaid D, Hong S, Marches R, Chen G, Bolisetty M, Baisch J, Walters L, Punaro M, Rossi RJ, Chung C-H, et al. Mapping systemic lupus erythematosus heterogeneity at the single-cell level. Nature Immunology (2020) 21:1094–1106. doi:10.1038/s41590-020-0743-0

26. Stoeckius M, Hafemeister C, Stephenson W, Houck-Loomis B, Chattopadhyay PK, Swerdlow H, Satija R, Smibert P. Simultaneous epitope and transcriptome measurement in single cells. Nature Methods (2017) 14:865–868. doi:10.1038/nmeth.4380

27. Kang HM, Subramaniam M, Targ S, Nguyen M, Maliskova L, McCarthy E, Wan E, Wong S, Byrnes L, Lanata CM, et al. Multiplexed droplet single-cell RNA-sequencing using natural genetic variation. Nature Biotechnology (2018) 36:89–94. doi:10.1038/nbt.4042

28. Jagger AL, Evans HG, Walter GJ, Gullick NJ, Menon B, Ballantine LE, Gracie A, Magerus-Chatinet A, Tiemessen MM, Geissmann F, et al. FAS/FAS-L dependent killing of activated human monocytes and macrophages by CD4+CD25-responder T cells, but not CD4+CD25+ regulatory T cells. J Autoimmun (2012) 38:29–38. doi:10.1016/j.jaut.2011.11.015

29. Kaplan MJ, Ray D, Mo RR, Yung RL, Richardson BC. TRAIL (Apo2 ligand) and TWEAK (Apo3 ligand) mediate CD4+ T cell killing of antigen-presenting macrophages. J Immunol (2000) 164:2897–2904. doi:10.4049/jimmunol.164.6.2897

30. Morita R, Schmitt N, Bentebibel S-E, Ranganathan R, Bourdery L, Zurawski G, Foucat E, Dullaers M, Oh S, Sabzghabaei N, et al. Human Blood CXCR5+CD4+ T Cells Are Counterparts of T Follicular Cells and Contain Specific Subsets that Differentially Support Antibody Secretion. Immunity (2011) 34:108–121. doi:10.1016/j.immuni.2010.12.012

31. Kim CH, Rott L, Kunkel EJ, Genovese MC, Andrew DP, Wu L, Butcher EC. Rules of chemokine receptor association with T cell polarization in vivo. J Clin Invest (2001) 108:1331–1339.

32. Tian Y, Babor M, Lane J, Schulten V, Patil VS, Seumois G, Rosales SL, Fu Z, Picarda G, Burel J, et al. Unique phenotypes and clonal expansions of human CD4 effector memory T cells re-expressing CD45RA. Nature Communications (2017) 8:1473. doi:10.1038/s41467-017-01728-5

33. Park D, Kim HG, Kim M, Park T, Ha H-H, Lee DH, Park K-S, Park SJ, Lim HJ, Lee CH. Differences in the molecular signatures of mucosal-associated invariant T cells and conventional T cells. Scientific Reports (2019) 9:7094. doi:10.1038/s41598-019-43578-9

34. Street K, Risso D, Fletcher RB, Das D, Ngai J, Yosef N, Purdom E, Dudoit S. Slingshot: cell lineage and pseudotime inference for single-cell transcriptomics. BMC Genomics (2018) 19:477. doi:10.1186/s12864-018-4772-0

35. Mailliard RB, Alber SM, Shen H, Watkins SC, Kirkwood JM, Herberman RB, Kalinski P. IL-18-induced CD83+CCR7+ NK helper cells. J Exp Med (2005) 202:941–953. doi:10.1084/jem.20050128

36. Robertson MJ. Role of chemokines in the biology of natural killer cells. JLeukoc Biol (2002) 71:173–183.

37. Rabacal W, Pabbisetty SK, Hoek KL, Cendron D, Guo Y, Maseda D, Sebzda E. Transcription factor KLF2 regulates homeostatic NK cell proliferation and survival. PNAS (2016) 113:5370–5375. doi:10.1073/pnas.1521491113

38. Hao Y, O’Neill P, Naradikian MS, Scholz JL, Cancro MP. A B-cell subset uniquely responsive to innate stimuli accumulates in aged mice. Blood (2011) 118:1294–1304. doi:10.1182/blood-2011-01-330530

39. Rubtsova K, Rubtsov AV, Thurman JM, Mennona JM, Kappler JW, Marrack P. B cells expressing the transcription factor T-bet drive lupus-like autoimmunity. J Clin Invest (2017) 127:1392–1404. doi:10.1172/JCI91250

40. Zhang W, Zhang H, Liu S, Xia F, Kang Z, Zhang Y, Liu Y, Xiao H, Chen L, Huang C, et al. Excessive CD11c+Tbet+ B cells promote aberrant TFH differentiation and affinity-based germinal center selection in lupus. PNAS (2019) 116:18550–18560. doi:10.1073/pnas.1901340116

41. Jenks SA, Cashman KS, Zumaquero E, Marigorta UM, Patel AV, Wang X, Tomar D, Woodruff MC, Simon Z, Bugrovsky R, et al. Distinct Effector B Cells Induced by Unregulated Toll-like Receptor 7 Contribute to Pathogenic Responses in Systemic Lupus Erythematosus. Immunity (2018) 49:725–739.e6. doi:10.1016/j.immuni.2018.08.015

42. Yu G, Wang L-G, Han Y, He Q-Y. clusterProfiler: an R Package for Comparing Biological Themes Among Gene Clusters. OMICS (2012) 16:284–287. doi:10.1089/omi.2011.0118

43. Baillie JK, Arner E, Daub C, De Hoon M, Itoh M, Kawaji H, Lassmann T, Carninci P, Forrest ARR, Hayashizaki Y, et al. Analysis of the human monocyte-derived macrophage transcriptome and response to lipopolysaccharide provides new insights into genetic aetiology of inflammatory bowel disease. PLoS Genet (2017) 13: doi:10.1371/journal.pgen.1006641

44. Schinnerling K, García-González P, Aguillón JC. Gene Expression Profiling of Human Monocyte-derived Dendritic Cells - Searching for Molecular Regulators of Tolerogenicity. Front Immunol (2015) 6: doi:10.3389/fimmu.2015.00528

45. Crowe LAN, McLean M, Kitson SM, Melchor EG, Patommel K, Cao HM, Reilly JH, Leach WJ, Rooney BP, Spencer SJ, et al. S100A8 & S100A9: Alarmin mediated inflammation in tendinopathy. Scientific Reports (2019) 9:1463. doi:10.1038/s41598-018-37684-3

46. Maeda A, Kawamura T, Ueno T, Usui N, Eguchi H, Miyagawa S. The suppression of inflammatory macrophage-mediated cytotoxicity and proinflammatory cytokine production by transgenic expression of HLA-E. Transplant Immunology (2013) 29:76–81. doi:10.1016/j.trim.2013.08.001

47. Hennequart M, Pilotte L, Cane S, Hoffmann D, Stroobant V, Plaen ED, Van den Eynde B. Constitutive IDO1 Expression in Human Tumors Is Driven by Cyclooxygenase-2 and Mediates Intrinsic Immune Resistance. Cancer Immunol Res (2017) 5:695–709. doi:10.1158/2326-6066.CIR-16-0400

48. Hamilton RG, Adkinson NF. In vitro assays for the diagnosis of IgE-mediated disorders. Journal of Allergy and Clinical Immunology (2004) 114:213–225. doi:10.1016/j.jaci.2004.06.046

49. Bendall SC. Diamonds in the doublets. Nature Biotechnology (2020) 38:559–561. doi:10.1038/s41587-020-0511-6

50. Giladi A, Cohen M, Medaglia C, Baran Y, Li B, Zada M, Bost P, Blecher-Gonen R, Salame T-M, Mayer JU, et al. Dissecting cellular crosstalk by sequencing physically interacting cells. Nature Biotechnology (2020) 38:629–637. doi:10.1038/s41587-020-0442-2

51. Cho BK, Wang C, Sugawa S, Eisen HN, Chen J. Functional differences between memory and naive CD8 T cells. PNAS (1999) 96:2976–2981. doi:10.1073/pnas.96.6.2976

52. Piqueras B, Connolly J, Freitas H, Palucka AK, Banchereau J. Upon viral exposure, myeloid and plasmacytoid dendritic cells produce 3 waves of distinct chemokines to recruit immune effectors. Blood (2006) 107:2613–2618. doi:10.1182/blood-2005-07-2965

53. Qiao Y, Giannopoulou EG, Chan CH, Park S, Gong S, Chen J, Hu X, Elemento O, Ivashkiv LB. Synergistic Activation of Inflammatory Cytokine Genes by Interferon-γ-Induced Chromatin Remodeling and Toll-like Receptor Signaling. Immunity (2013) 39:454–469. doi:10.1016/j.immuni.2013.08.009

54. Baillie JK, Arner E, Daub C, De Hoon M, Itoh M, Kawaji H, Lassmann T, Carninci P, Forrest ARR, Hayashizaki Y, et al. Analysis of the human monocyte-derived macrophage transcriptome and response to lipopolysaccharide provides new insights into genetic aetiology of inflammatory bowel disease. PLoS Genet (2017) 13: doi:10.1371/journal.pgen.1006641

55. Stoeckius M, Zheng S, Houck-Loomis B, Hao S, Yeung BZ, Mauck WM, Smibert P, Satija R. Cell Hashing with barcoded antibodies enables multiplexing and doublet detection for single cell genomics. Genome Biology (2018) 19:224. doi:10.1186/s13059-018-1603-1

56. Lawlor N, Márquez EJ, Orchard P, Narisu N, Shamim MS, Thibodeau A, Varshney A, Kursawe R, Erdos MR, Kanke M, et al. Multiomic Profiling Identifies cis-Regulatory Networks Underlying Human Pancreatic β Cell Identity and Function. Cell Rep (2019) 26:788–801.e6. doi:10.1016/j.celrep.2018.12.083

57. Varshney A, Scott LJ, Welch RP, Erdos MR, Chines PS, Narisu N, Albanus RD, Orchard P, Wolford BN, Kursawe R, et al. Genetic regulatory signatures underlying islet gene expression and type 2 diabetes. Proc Natl Acad Sci USA (2017) 114:2301–2306. doi:10.1073/pnas.1621192114

58. Scrucca L, Fop M, Murphy TB, Raftery AE. mclust 5: Clustering, Classification and Density Estimation Using Gaussian Finite Mixture Models. R J (2016) 8:289–317.

59. Butler A, Hoffman P, Smibert P, Papalexi E, Satija R. Integrating single-cell transcriptomic data across different conditions, technologies, and species. Nature Biotechnology (2018) 36:411–420. doi:10.1038/nbt.4096

60. Blondel VD, Guillaume J-L, Lambiotte R, Lefebvre E. Fast unfolding of communities in large networks. J Stat Mech (2008) 2008:P10008. doi:10.1088/1742-5468/2008/10/P10008

61. Saelens W, Cannoodt R, Todorov H, Saeys Y. A comparison of single-cell trajectory inference methods. Nature Biotechnology (2019) 37:547–554. doi:10.1038/s41587-019-0071-9

