## Supplementary Figures for "Single cell analysis of blood mononuclear cells stimulated through CD3 and CD28 shows collateral activation of B and NK cells and demise of monocytes"

**Fig. S1: De-multiplexing of singlet cells using cell hashtag oligonucleotides (HTOs) and individual genetic variation.** (A) Distributions of HTO normalized read counts across all PBMCs were bimodal for each condition. Two component gaussian mixture models were fit to each distribution to classify each cell as positive/negative for each HTO. (B) Decision tree for singlet and multiplet cell characterization. HTOs were first used to remove multiplets from 2 or more conditions, while *Demuxlet* and individual genotypes were used to remove multiplets from 2 or more individuals. Lastly, cell-specific surface protein expression was used to identify and remove heterotypic cell multiplets. (C) After de-multiplexing HTO information, the numbers of singlet cells (positive for only one HTO), multiplet cells (positive for two or more HTOs), and empty droplets (positive for no HTOs). (D) After de-multiplexing individual/genotype information, the numbers of singlet (SNG; corresponding to a single donor), doublet (DBL; corresponding to two or more donors), and ambiguous (AMB; unable to assign to any donor) droplets. (E) The overlap of cell droplet classifications after de-multiplexing with HTOs and donor genotypes. Empty droplets (positive for no HTOs) were commonly annotated as ambiguous (unable to assign to any donor via *Demuxlet*), while doublet/triplet (positive for two or more HTOs) were annotated more commonly as DBL via *Demuxlet*. (F) Comparison of donor proportions of CD69^+^/CD25^+^ PBMCs at baseline and anti-CD3/CD28 stimulated conditions in flow cytometry and ADT (CITE-seq) assays. R represents the Pearson’s correlation coefficient. Note the high correlation between cell compositions from flow and CITE-seq data.

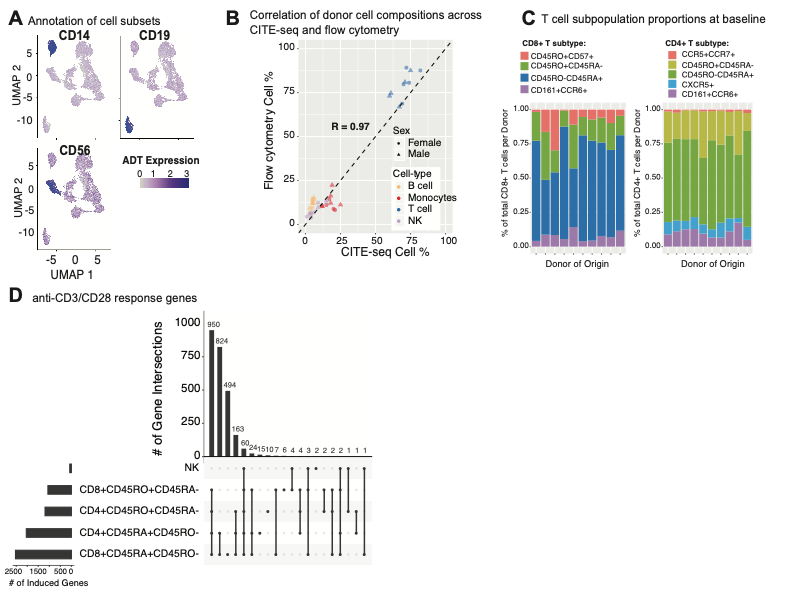

**Fig. S2: Characterizing T cell responses to anti-CD3/CD28 stimulation.** (A) Annotation of monocytes (CD14), B cells (CD19), and NK cells (CD56) using ADT expression of cell surface markers. (B) Donor PBMC cell-type proportions from flow cytometry and CITE-seq experiments were positively correlated. R = Pearson correlation coefficient. (C) Proportions of (left) CD8^+^ and (right) CD4^+^ T cell subpopulations across each of the 10 donors. (D) Upset diagram depicting the common and unique anti-CD3/CD28 response genes in NK and T cells.

**Fig. S3: LPS stimulation induces an inflammatory response in monocytes, but not in other PBMC subsets.** (A) Annotation of PBMC cell subsets using cell surface protein expression. (B) LPS-activated monocytes show more than 2-fold higher expression of pro-inflammatory transcripts (e.g., *IL8, IL-1B*). (C) Pathway enrichment analysis (using Gene ontology biological process terms) identifies processes associated with genes induced/reduced in LPS-activated monocytes respectively. FDR = false discovery rate; GeneRatio = proportion of overlap of response genes with the pathway associated genes (D) Distribution of monocytes in the inferred pseudotime trajectory reveal cells at baseline (blue) are near the trajectory start, while LPS-activated cells (red) are positioned towards the trajectory end. (E) Top genes and proteins whose expression is correlated with pseudotime. Expression in the heatmap is represented as z-scored values. (F) Pseudotime states of baseline and LPS-activated monocytes are comparable across the 10 PBMC donors.

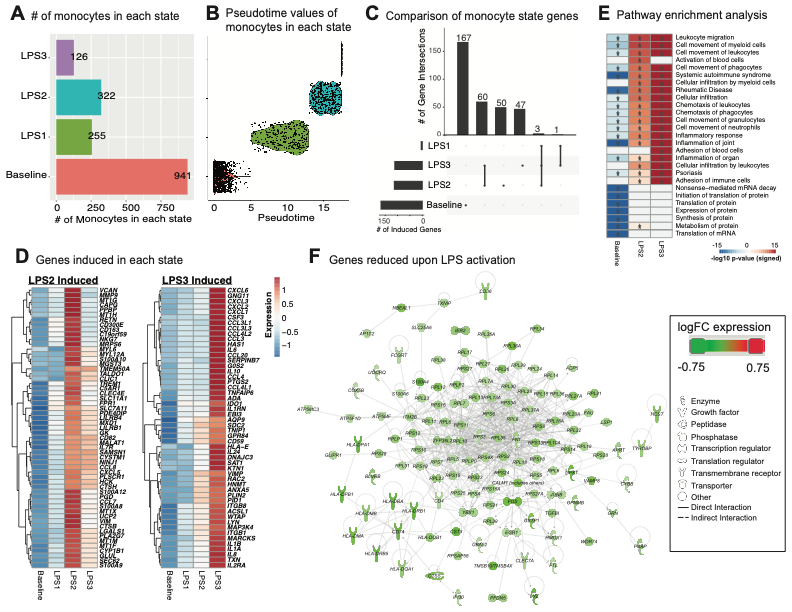

**Fig. S4: Investigating the heterogeneity of monocyte responses to LPS.** (A) Numbers of monocytes found in each of the four states. (B) Monocytes in each cluster had mutually exclusive pseudotime values. (C) Upset diagram detailing the intersections of genes induced in each monocyte state. Induced genes were defined by comparing the average expression of the gene in that state vs. the average expression of the gene in all other states combined (e.g., Baseline vs. all LPS1, LPS2, LPS3 states). (D) Heatmaps showing the average z-scored expression values of the top genes induced in LPS2 and LPS3 respectively. (E) Pathway analysis of IPA diseases and biological function terms for genes induced at baseline, LPS2, and LPS3 states. Asterisks in the heatmap “*” signify a p-value < 0.05. (F) Network of the genes reduced in monocytes upon LPS activation Molecules are colored based on their log2 fold change in expression of all activated monocyte states vs. those at baseline.
